# Serum IgM crosslinks hepatitis B subviral particles into supramolecular immune complexes

**DOI:** 10.64898/2026.09.30.755772

**Authors:** Chang Liu, Aditi Dhawan, Xinyi Yao, Juyeon Park, Gautam Penna, Sean Mulligan, Alexandra Beaver, Helen Chen, Flora Donati, Stephan Urban, Ralf Bartenschlager, George Georgiou, Shirin Nkongolo, Stefan Seitz, Zunlong Ke

## Abstract

Hepatitis B virus (HBV)-infected cells release hepatitis B surface antigen (HBsAg)-rich subviral particles (SVPs) in large excess over infectious virions. SVPs bind antibodies against HBsAg (anti-HBs), including neutralizing antibodies, but the physical products of antibody capture in human blood are uncharacterized. Here we used cryo-electron tomography (cryo-ET) to image HBsAg-enriched serum from a patient with severe acute Hepatitis B and observed three-dimensional assemblies of intact IgM-SVP immune complexes. These associations persisted at a lower particle density and were validated by a spatial-randomization null model. Subtomogram averaging and single-particle cryo-electron microscopy (cryo-EM) identified three serum IgM Fc-core assemblies, IgM-J-CD5L, IgM-J and IgM-J-SC, with cryo-EM maps reaching 2.8 Å resolution. Mapping these Fc core assemblies into tomograms revealed full-length IgM engaging individual SVPs through several Fab arms and bridging neighboring particles into immune complex networks containing multiple IgM molecules and SVPs. Thus, serum IgM uses multivalent Fab interactions to crosslink HBsAg-rich SVPs into supramolecular immune complexes. This observation extends the SVP decoy model from passive antibody sequestration to higher-order particle organization.

## Introduction

Hepatitis B virus (HBV) remains a major cause of acute and chronic liver disease worldwide^1,2^. Infected hepatocytes release not only infectious Dane particles but also excessive numbers of non-infectious spherical and filamentous subviral particles (SVPs), a characteristic feature of HBV infection^3^; these SVPs are composed largely of hepatitis B surface antigen (HBsAg)^4^. Because SVPs display the same surface antigens as virions, they can intercept antibodies against HBsAg (anti-HBs); purified SVPs reduce antibody-mediated neutralization of infectious HBV in vitro^5^. This immune-decoy model explains why excess particles could protect virions, but it leaves a basic physical question unanswered: what structures are created when native, full-length antibodies encounter repetitive HBsAg on SVPs in human blood?

HBsAg-IgM immune complexes have been detected in patient sera for decades. Early biochemical studies showed that HBsAg co-fractionates with IgM in patient serum^6,7^. HBsAg- and immunoglobulin-containing circulating immune complexes are also common during acute hepatitis B and can be captured by C1q-based assays^8^. These studies establish that HBsAg and IgM coexist in circulating immune complexes capable of binding C1q, but bulk assays did not reveal how many particles are connected, how Fab arms are deployed, or how molecular IgM composition relates to higher-order architecture. HBsAg-IgM complexes have also been detected during chronic HBV infection^7,9^, and IgG anti-HBs has been identified in circulating immune complexes during both acute and chronic infection^10^.

These architectural properties are biologically consequential because connectivity changes both the effective valency and physical scale of immune complexes. For IgM, antigen-induced Fc organization creates a platform for C1q engagement and complement activation^11,12^. More broadly, immune-complex size can influence clearance kinetics^13^. Determining whether IgM simply coats individual SVPs or crosslinks them into higher-order assemblies is therefore essential for understanding the physical consequences of the HBV decoy strategy.

IgM is particularly suited to translate recognition of repetitive antigens into supramolecular assembly. Pentameric IgM presents ten Fab arms around an asymmetric Fc core^14^. The core contains the J chain and can additionally accommodate CD5L or secretory component (SC), generating compositionally distinct IgM assemblies^15^. Structural studies of purified IgM, including recombinant complexes and myeloma-derived pentamers, have defined IgM-J, IgM-J-CD5L and IgM-J-SC Fc cores^16–20^. Complementing these structures, serum proteomics identified CD5L as a canonical component of native circulating IgM^21,22^. In parallel, cryo-EM studies have defined the architectures of native and recombinant SVPs^23–25^ and the molecular interfaces between HBsAg and antibody Fabs^26,27^. Although these individual structures are increasingly well defined, the supramolecular organization that emerges when full-length serum IgM engages patient-derived SVPs remains unresolved.

Here we use a structural serology strategy integrating cryo-electron tomography (cryo-ET), single-particle cryo-electron microscopy (cryo-EM), and spatial modelling to define IgM-SVP complexes in HBsAg-enriched serum fractions from patients with HBV infection^28,29^. We directly visualize intact IgM-SVP assemblies and show, using a spatial-randomization null model, that these associations exceed random proximity. We identify three Fc-core assemblies of native serum IgM and show that full-length IgM uses multivalent Fab interactions to crosslink SVPs into supramolecular immune complexes, extending the SVP decoy model beyond passive antibody sequestration.

## Results

### Cryo-ET reveals native IgM-SVP complexes

To visualize native IgM-SVP complexes in patient serum, we prepared HBsAg-enriched serum fractions from two individuals infected with HBV genotype D1: one with severe acute hepatitis B and liver failure and the other with chronic infection and limited biochemical liver injury (Supplementary Table 1). HBV-containing material was enriched by sequential ultracentrifugation and chromatographic fractionation, guided by HBsAg and HBV DNA measurements, using our previously established method for purifying serum-derived HBV particles (Supplementary Fig. 1)^30^. The severe acute sample was used for the IgM-SVP tomography and the principal structural analyses, whereas the chronic sample provided an independent comparison of native serum IgM Fc-core architecture.

Cryo-ET directly revealed the three-dimensional organization of native IgM-SVP assemblies in the severe acute sample. We collected 269 tilt series and retained 253 tomograms after quality control. High-concentration tomograms contained numerous spherical SVPs together with intact IgM molecules (Fig. 1a and Supplementary Movie 1). The central Fc region and flexible Fab arms of individual IgM molecules were distinguishable in tomographic slices and 3D segmentations. Enlarged views showed IgM molecules adjacent to one or more SVPs, with Fab-shaped densities extending toward the particle surfaces (Fig. 1b-d).

**Fig. 1.**
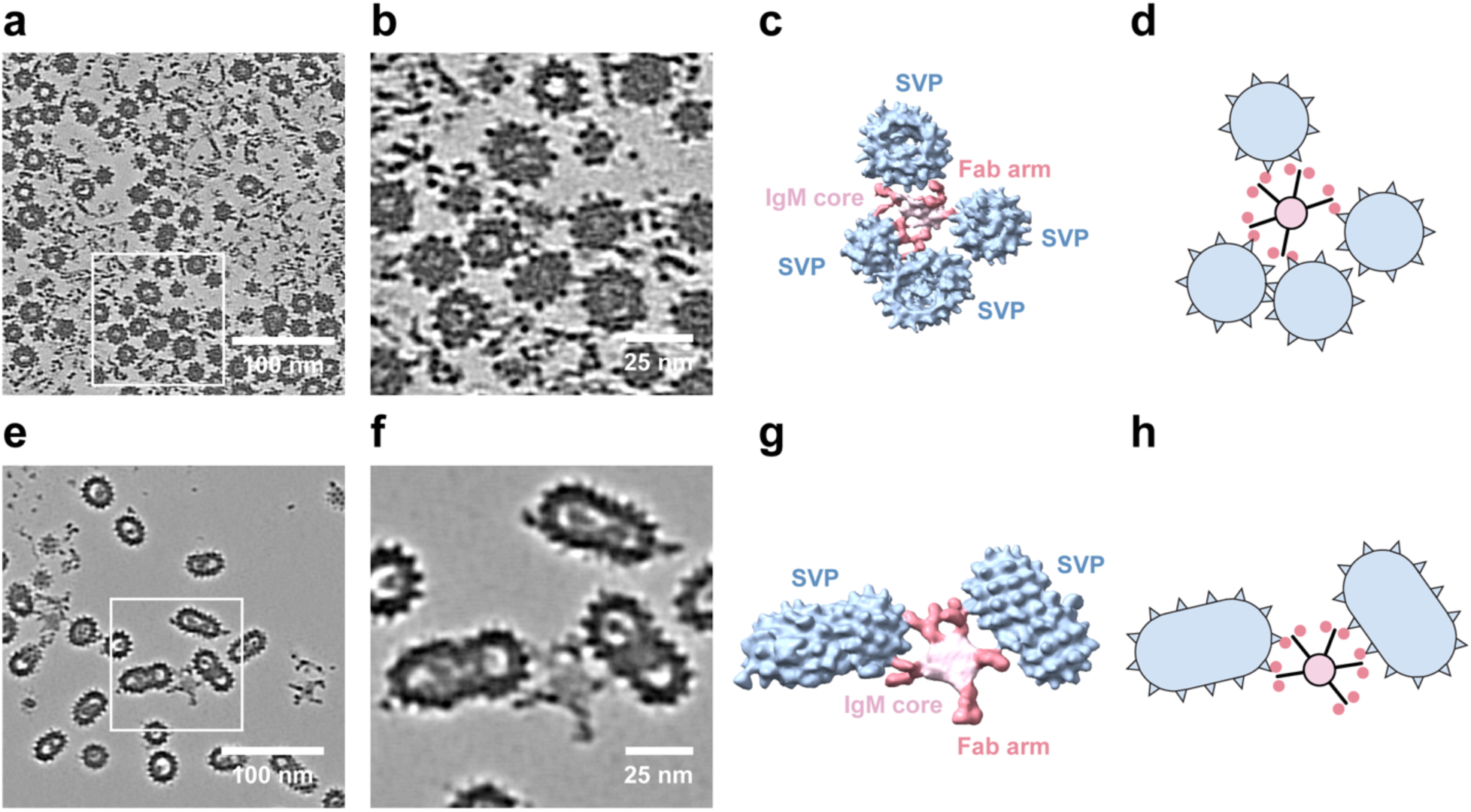
Cryo-ET visualization of native IgM-SVP complexes. **a-d,** Representative tomographic slice (**a**), enlarged view of the boxed region (**b**), corresponding three-dimensional segmentation (**c**), and schematic representation of the three-dimensional segmentation (**d**) from a high-concentration HBsAg-enriched serum fraction from the patient with severe acute hepatitis B. Spherical SVPs and intact IgM molecules are visible, with Fab-shaped densities extending toward the surfaces of one or more neighboring SVPs. **e-h,** Representative tomographic slice (**e**), enlarged view of the box region (**f**), corresponding three-dimensional segmentation (**g**), and schematic representation of the three-dimensional segmentation (**h**) and from a neighboring, lower-concentration HBsAg-enriched fraction. Isolated IgM-SVP associations remain visible under the lower particle density. Segmentation colors: SVPs, light blue; IgM Fc regions, light pink; Fab arms, salmon.

To determine whether IgM molecules positioned adjacent to one or more SVPs reflected genuine association rather than incidental proximity arising from high particle density, we imaged a neighboring, lower-concentration fraction from the same HBsAg-enriched preparation. This fraction contained substantially fewer densities of both IgM and SVPs, yet isolated IgM-SVP associations remained clearly visible in raw tomographic slices and three-dimensional renderings (Fig. 1e-h and Supplementary Movie 2). Because the lower particle density in this fraction reduces the likelihood of incidental neighbors while preserving true interactions, these sparse tomograms provided the appropriate dataset for the spatial-randomization null model applied below.

We also observed smaller, Y-shaped antibody densities associated with SVPs, consistent with IgG (Supplementary Fig. 2a). Occasional examples showed a single IgG-like molecule bridging two SVPs, with one Fab arm contacting each particle (Supplementary Fig. 2b). IgG-like binding events were less frequently identifiable in the imaged fractions, and larger assemblies containing multiple IgG-like molecules and SVPs were rarely observed. The limited number of identifiable IgG-SVP complexes did not support structural and quantitative analyses comparable to those of IgM. We therefore focused on the recurrent, readily recognizable pentameric IgM assemblies, while retaining the IgG observations as evidence that SVP binding and interparticle bridging are not restricted to IgM.

### Cryo-EM resolves native serum IgM Fc-core assemblies

Subtomogram averaging and 3D classification of the higher-concentration cryo-ET dataset resolved heterogeneity within the IgM Fc core and resolved three classes at 7.6-11.4 Å resolution (Supplementary Fig. 3). Distinct densities attributable to CD5L and SC indicated that compositionally different IgM assemblies coexist within the same serum-derived fractions that contain the visualized IgM-SVP complexes. To define these states at higher resolution, we applied single-particle cryo-EM to the same severe acute serum fraction, which resolved the same three assemblies at near-atomic resolution: IgM-J-CD5L (53.5%, 2.8 Å), IgM-J (25.0%, 3.1 Å) and IgM-J-SC (21.5%, 3.0 Å) (Fig. 2a-c and Supplementary Fig. 4), with side-chain and glycan density resolved across much of each Fc core (Fig. 2d-e and Supplementary Figs. 4-5). Because cryo-EM and cryo-ET image processing workflows differ in particle selection and classification strategy, their class proportions are not estimates of absolute serum abundance.

**Fig. 2.**
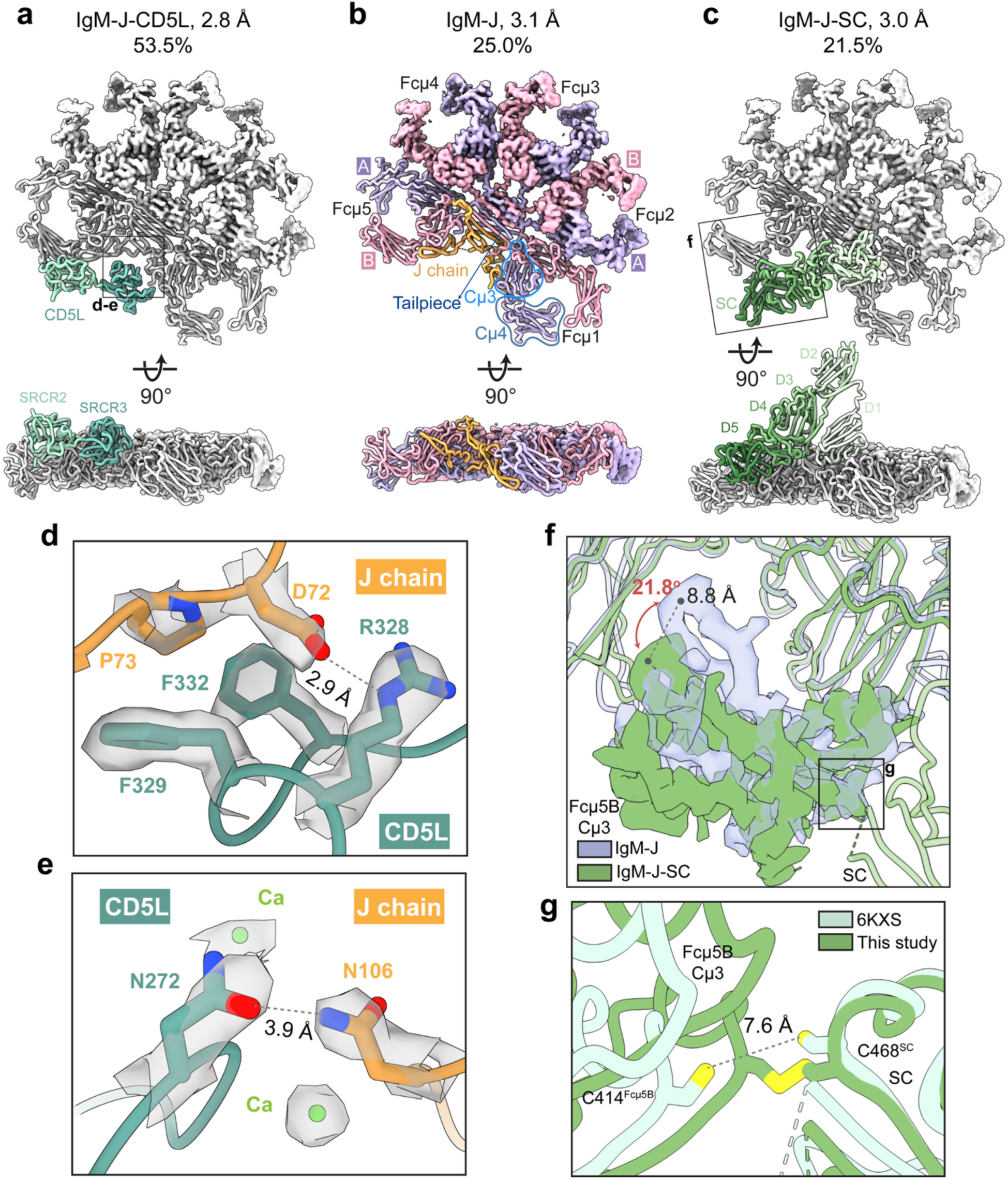
Cryo-EM structures of native serum IgM Fc-core assemblies. **a-c,** Cryo-EM structures of the three native serum IgM Fc-core assemblies, IgM-J-CD5L (**a**), IgM-J (**b**) and IgM-J-SC (**c**), shown as top and side views (90° rotation), with corresponding resolutions and particle-class distributions. The IgM pentamer is colored in alternating light pink and lavender to distinguish individual Fcμ protomers; CD5L is colored teal, SC is colored green, and the J chain is colored orange. Boxed regions indicate the areas magnified in **d-f**. **d,** Representative cryo-EM density at the CD5L-J chain interface, illustrating the well-resolved side-chain densities that support atomic modeling. **e,** Newly resolved interaction between N272^CD5L^ and N106^J^. Side-chain densities are shown. **f,** Comparison of the IgM-J and IgM-J-SC structures from the same patient sample, showing a ∼21.8° swing and ∼8.8 Å distal displacement of the Fcμ5B Cμ3 domain upon SC binding. **g,** Comparison of IgM-J-SC structures, showing the C468^SC^-C414^Fcμ5B^ disulfide bond observed in our structure but not in PDB 6KXS, in which the two cysteine sulfur atoms are separated by ∼7.6 Å.

The native Fc-core structures reproduce the overall architectures reported for recombinant and purified IgM complexes (Fig. 2d, Supplementary Fig. 6, and Supplementary Movie 3)^16–18,20^. CD5L occupies the asymmetric cavity and interacts predominantly with the J chain. Within this interface, our maps resolve a CD5L-J-chain contact, N272^CD5L^-N106^J^, located adjacent to the Ca^2+^-binding pocket. This contact is weak or absent in previously deposited models (Fig. 2e and Supplementary Fig. 7a)^18,20^, extending the molecular description of CD5L engagement with the J chain.

SC contacts both the J chain and Fc core. Relative to IgM-J, SC association displaces the Fcμ5B Cμ3 domain by ∼22°, shifting its distal region by ∼8.8 Å (Fig. 2f). Beyond this global rearrangement, SC-associated contacts, including T33^SC^-D131^J^, S51^SC^-E468^Fcμ1B^ and E331^SC^-Y576^Fcμ5A^ (Supplementary Fig. 7b-d), extend the structural description of partner recognition without implying altered binding affinity. In addition, our serum IgM structure indicates that C468^SC^ forms a disulfide bond with C414^Fcμ5B^, consistent with PDB 7K0C^17^ but not in PDB 6KXS^16^, with the later study’s corresponding sulfur atoms are separated by ∼7.6 Å (Fig. 2g and Supplementary Fig. 7d). Although sample preparation-dependent effects cannot be excluded, this covalent linkage provides an additional point of stabilization within the IgM-J-SC assembly^16^.

To determine whether the three Fc-core assemblies identified in the severe acute sample reflect a general feature of serum IgM rather than a property of a single patient, we applied the same cryo-EM analysis to the chronic-infection sample. This independent dataset reproduced IgM-J-CD5L (64.5%; 4.0 Å) and IgM-J (35.5%; 5.9 Å), but yielded no IgM-J-SC class (Supplementary Fig. 8), indicating that the two major assemblies are consistent across patients whereas SC association is variable. To assess this variability more broadly, we used capture ELISA to measure total, CD5L-associated and SC-associated IgM in serum from healthy donors (n = 4) and HBV patients (n = 6) (Supplementary Fig. 9 and Supplementary Table 2). Relative to healthy donors, SC-associated IgM was elevated in three of six HBV patients. Circulating SC and secretory IgM have been linked to liver disease and severe hepatobiliary injury, but their levels vary among patients with chronic liver disease^31–33^, a plausible source of this variability. Whether this variability reflects hepatobiliary transport, altered clearance, or other aspects of liver disease remains to be determined, and our data do not establish an association between IgM-J-SC and a specific stage of HBV infection.

### IgM-SVP engagement is non-random and multivalent

To further test whether the IgM-SVP associations visualized by cryo-ET reflect genuine engagement rather than incidental co-occurrence, we applied a Monte Carlo simulation to randomize the spatial localization of IgM and SVPs on the low-concentration dataset described above (Fig. 3a-b). For each tomogram, we generated randomized particle locations and orientations that preserved the number of IgM and SVP particles and their measured sizes, while considering the excluded volume and the accessible tomographic volume and boundaries (Supplementary Fig. 10a-c), then compared the frequency of possible IgM-SVP associations given the SVP’s nearest neighbor to IgM distances between the observed and randomized configurations. Across 3 tomograms (observed) and 100 randomized realizations per tomogram (simulation), we observed a sharp peak around the interaction cutoff distance (33.1 nm) in the real tomograms, but not in the simulated ones, indicating that IgM-SVP associations are enriched relative to chance (Fig. 3c, Supplementary Fig. 10d-g). To further assess the number of SVPs per IgM interaction, we plotted the histogram based on the number of SVPs per IgM interacted in the real and simulated tomograms (Supplementary Fig. 10h). The results show the observed associated frequency was 1.1 SVPs/IgM, compared with 0.32 SVPs/IgM with a 95% confidence interval between 0.16 and 0.48, and sigma of 0.08. Next, we asked whether there is difference between the abundance of IgM binds to two or more SVPs (multivalency) in the real and simulated tomogram. Our simulation indicates that there are statistically significantly more IgMs with two or more SVPs bound in the real data (p< 0.0001), indicating the multivalency and possible network is non-random (Fig. 3d).

**Fig. 3.**
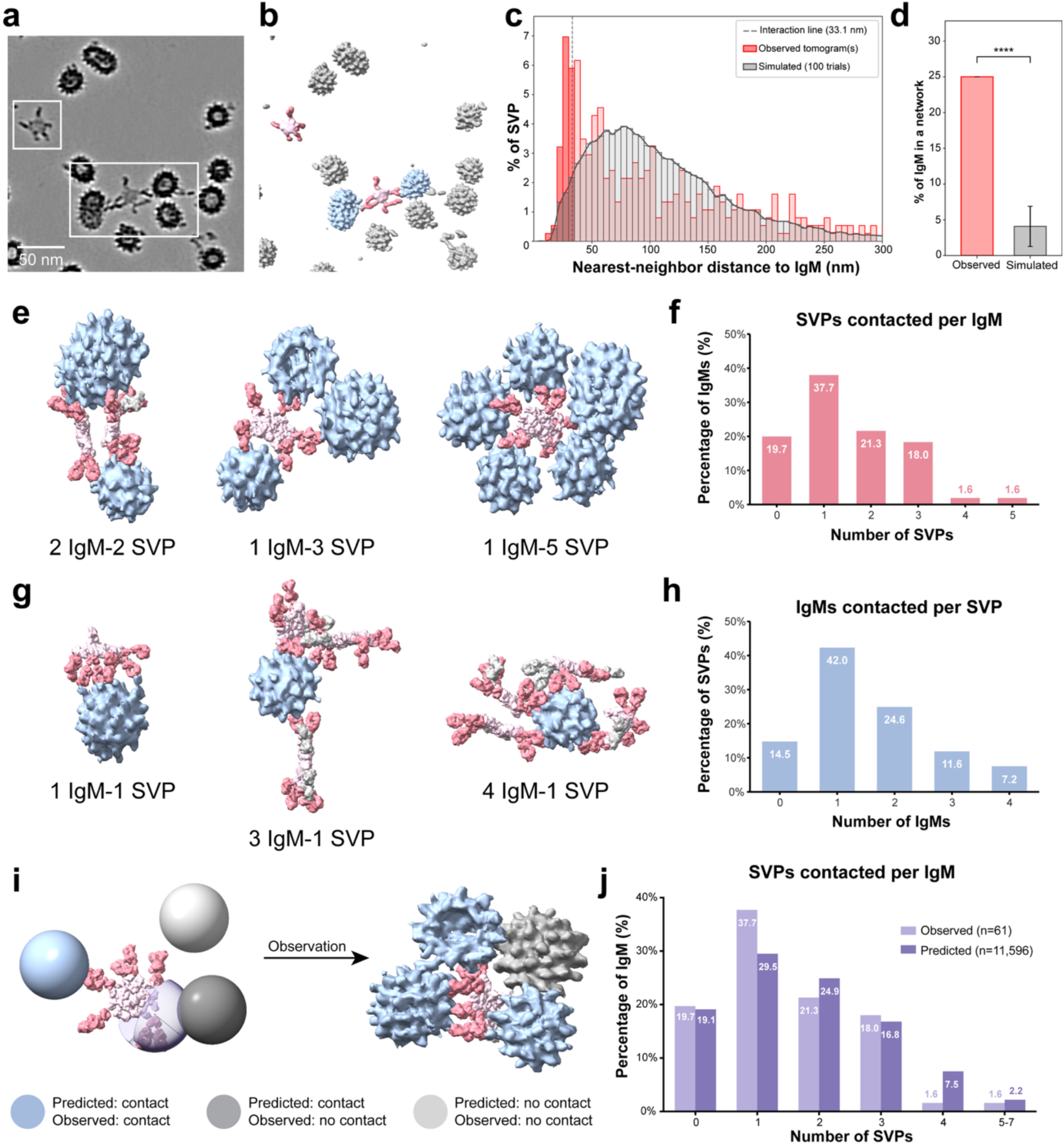
IgM-SVP complexes engage non-randomly and multivalently. **a-d,** Spatial-randomization null test. **a,** Representative tomographic slice from a low-concentration HBsAg-enriched fraction, showing free and IgM-associated SVPs together with free and SVP-associated IgM. **b,** Corresponding three-dimensional segmentation of the particles in **a**. **c,** Distribution of nearest-neighbor distances from SVPs to IgM molecules in the observed tomogram compared with distributions obtained from 100 spatial-randomization simulations. The dashed line indicates the interaction-distance threshold (33.1 nm) **d,** Percentage of IgM molecules participating in interaction networks containing at least two contacted SVPs in the observed tomogram compared with spatial-randomization simulations. (P = 1,13×10^−13^, n=100). **e,f,** IgM-centered analysis of multivalent SVP engagement. **e,** Representative renderings of individual IgM molecules engaging one to five SVPs. **f,** Observed frequency distribution of the number of SVPs bound per IgM molecule (n = 61 IgM molecules). **g,h,** SVP-centered analysis of multivalent IgM engagement. **g,** Representative renderings of individual SVPs engaged by one to four IgM molecules. **h,** Observed frequency distribution of the number of IgM molecules bound per SVP (n = 69 SVPs). **i,j,** Geometry-based prediction of IgM-SVP multivalent engagement. **i,** Schematic of the geometric analysis used to predict expected SVP-IgM engagement frequencies. **j,** Geometry-based predicted versus observed frequency distributions of SVPs bound per IgM, as in **f**. Segmentation colors: SVPs, light blue; IgM Fc regions, light pink; Fab arms, salmon.

To quantify IgM-SVP multivalency, we first reconstructed full-length IgM molecules, including their Fc cores and associated Fab arms. We used the higher-concentration tomograms for this analysis because the high particle numbers facilitate subtomogram alignment, orientation determination and Fc-core classification. Because Fab arms are generally too flexible to resolve by subtomogram averaging, we manually segmented individual Fab densities in the tomograms, using the Fc cores resolved above as positional references. This reconstruction revealed two forms of multivalent engagement: Fab arms from one IgM extending toward separate neighboring SVPs (interparticle bridging), and several Fab arms from one IgM contacting the same SVP (intraparticle multivalency) (Fig. 3e-h and Supplementary Figs. 11-12).

Manual inspection of 61 IgM molecules and their neighboring SVPs in the higher-concentration tomograms quantified this multivalency directly: individual IgM molecules engaged as many as five SVPs (Fig. 3e), with the majority engaging one SVP (37.3%), followed by two (25.3%) and three (15.3%) SVPs, whereas engagement of four or five SVPs was rare (1.6% each); 18.7% of IgM molecules showed no SVP engagement (Fig. 3f). The reciprocal relationship also held: individual SVPs were engaged by as many as four IgM molecules (Fig. 3g), with 42.3% contacted by one IgM, 24.3% by two, 11.6% by three, and 7.2% by four, while 14.6% showed no IgM engagement (Fig. 3h). Similar multivalent engagement was also observed in the low-concentration tomograms, indicating that these interactions are not an artifact of particle crowding (Supplementary Fig. 13 and Supplementary Movie 2).

To extend this quantification beyond manually inspected particles, we developed a geometric Fab-reach model that predicts SVP contacts from the measured IgM Fc-core position and orientation, the Fab hinge location and the experimentally sampled Fab angular range, up to ±60° in-plane rotation, ±110° out-of-plane tilt and ±180° spin (Fig. 3i, Supplementary Fig. 14). Because this prediction also depends on the size and position of each neighboring SVP, we classified the pleomorphic SVP population by subtomogram averaging, separating 61,963 particles from 253 tomograms into seven diameter classes ranging from 22 to 28 nm (Supplementary Fig. 15). Each particle’s measured class, size and position were used in the model rather than approximating SVPs as a single idealized sphere. Applied across 133 tomograms containing 11,569 IgM molecules and 26,816 SVPs after quality control, the model predicted that one- and two-SVP contacts were the predominant engagement modes, each involving approximately one quarter of IgM molecules, while fewer than 10% reached four or more SVPs, closely matching the manually quantified frequencies described above (Fig. 3j and Supplementary Fig. 16). This distribution held across IgM-J-CD5L, IgM-J and IgM-J-SC populations (Fig. 2a-c and Supplementary Fig. 3), indicating that multivalent SVP engagement is a shared feature of serum IgM rather than restricted to a specific Fc-core composition.

### IgM crosslinks SVPs into supramolecular immune complexes

Bidirectional multivalency implies that IgM-SVP contacts need not remain local, isolated events. We therefore searched for higher-order, supramolecular assemblies extending beyond simple pairwise engagement. Using the geometric prediction described above, we traced a larger interaction network and identified a candidate region of extensive connectivity. To confirm and visualize this assembly accurately, we manually traced the Fab densities and segmented the SVPs within this region, revealing a cluster of 10 IgM molecules and 10 SVPs, all mutually connected into a continuous network spanning approximately 200 nm by 100 nm, in which IgM bridges SVPs into a single interconnected immune complex (Fig. 4a-b). A smaller-scale higher-order assembly, comprising 2-3 IgM molecules and 3-5 SVPs, was also identified in the low-concentration tomograms (Supplementary Fig. 13c-d), indicating that this crosslinked architecture is not a consequence of particle crowding at higher concentration.

**Fig. 4.**
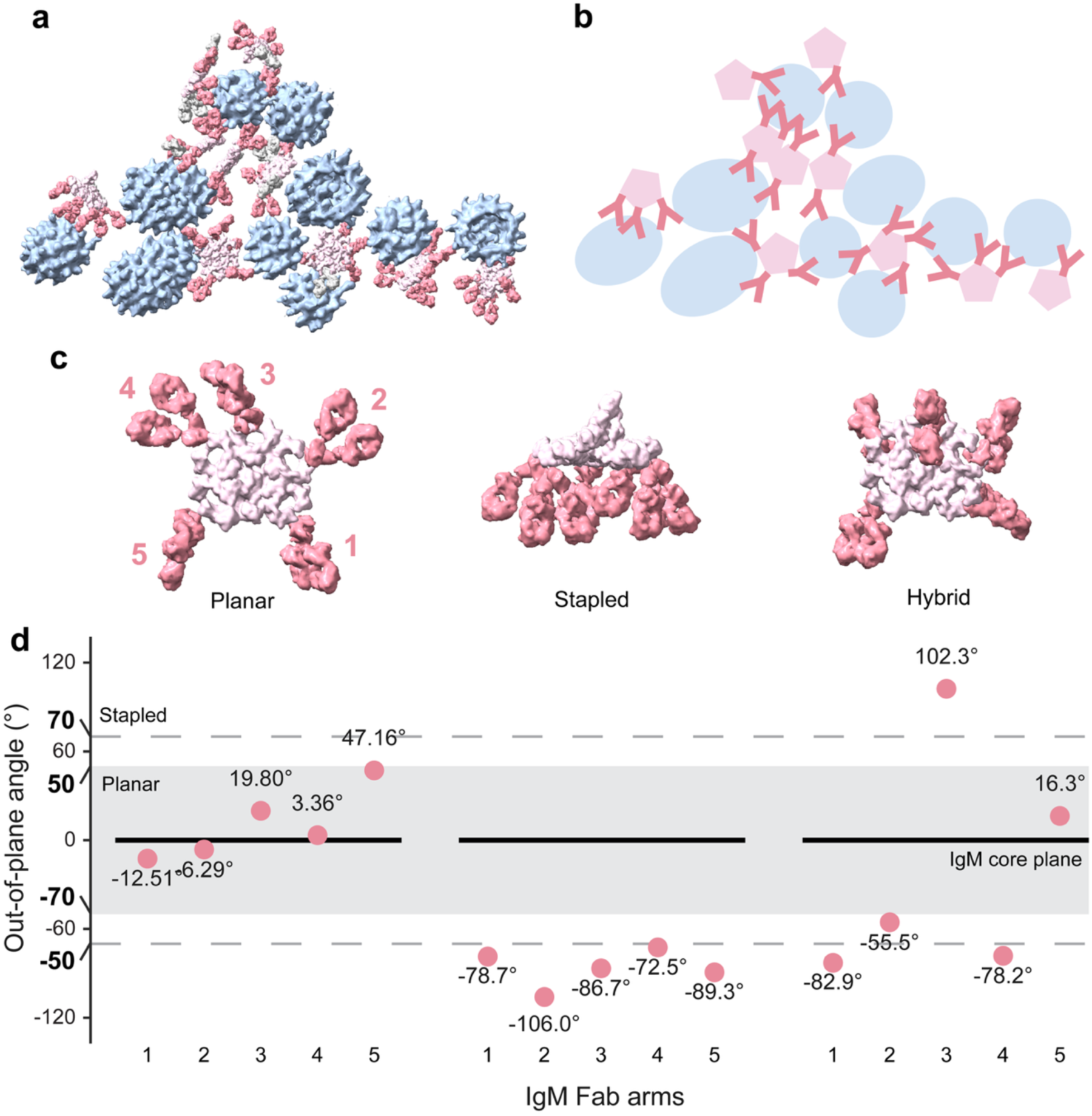
IgM crosslinks SVPs into supramolecular assemblies. **a-b,** Higher-order immune complex of IgM molecules and SVPs. **a,** Three-dimensional segmentation of an interconnected network comprising 10 IgM molecules and 10 SVPs. **b,** Cartoon representation of **a**. **c,** Full-length IgM architecture when bound to SVPs, shown as IgM-only segmentations in planar (left), stapled (middle) and hybrid (right) conformations. **d,** Out-of-plane Fab arm angles for the planar (left), stapled (middle) and hybrid (right) conformations shown in **d-f**, plotted across the five Fab arm positions. Planar IgM has all Fab arms within ±20°; stapled IgM has all Fab arms beyond ±50° of the same sign; hybrid IgM contains arms of both signs, with at least one arm meeting the planar or stapled criterion. Segmentation colors: SVPs, light blue; IgM Fc regions, light pink; Fab arms, salmon.

Within these networks, full-length IgM adopted one of three Fab-arm conformations (Fig. 4c): planar, in which Fab arms extend away from the Fc core, and stapled, in which Fab arms bend toward one side of the Fc core, both consistent with classical descriptions of IgM architecture^34^; and a third, hybrid conformation, in which some arms extend while others bend. We classified these conformations quantitatively from the out-of-plane angle of each of the five Fab arms: planar IgM had all Fab arms within ±50°, stapled IgM had all Fab arms beyond ±70° while residing the same side of Fc core, and hybrid IgM is non-planar, non-stapled IgM class (Fig. 4d).

The stapled conformation clusters Fab arms on one side of the IgM molecule, a geometry compatible with C1q engagement and complement activation^11,12^; because C1q was not present in our structures, this comparison reflects geometric compatibility rather than evidence of activation. Stapled IgM was uncommon in both datasets, accounting for 3 of 43 SVP-bound IgM molecules in the higher-concentration tomograms and 2 of 18 isolated IgM molecules, indicating that although individual IgM-SVP complexes can adopt a C1q-compatible conformation, this configuration represents a minor fraction of the population.

Together, these observations indicate that serum IgM does not merely coat individual SVPs, as envisioned by a passive sequestration model in which IgM engages multiple SVPs or SVPs engage multiple IgM without interconnection, but instead crosslinks SVPs into interconnected, supramolecular immune complexes (Supplementary Fig. 17). This higher-order organization extends the SVP decoy model beyond passive antibody sequestration: rather than simply confirming that IgM-HBsAg complexes exist, as established biochemically for decades, our data reveal that these complexes assemble into larger, interconnected networks bridging multiple IgM and SVPs. Because immune-complex connectivity can influence clearance kinetics and complement engagement^11,13^, the transition from isolated to crosslinked assemblies may carry physiological consequences for how the immune system processes HBsAg-rich serum.

## Discussion

The stoichiometry and architecture of HBV SVP-IgM complexes in patient serum have remained undefined. We have used cryo-ET to visualize intact IgM-SVP assemblies from serum (Fig. 1a-d). Imaging a naturally lower-concentration serum fraction reduces crowding without eliminating the assemblies (Fig. 1e-h), and a spatial null model shows enrichment of SVP-IgM interactions beyond random proximity (Fig. 3a-d). These independent observations support antigen-associated immune complexes rather than co-localization alone.

Cryo-EM analyses resolved the molecular identity of the IgM engaged in these complexes. The severe acute sample contained three distinct Fc-core assemblies, IgM-J-CD5L, IgM-J and IgM-J-SC, while the chronic sample yielded the first two states (Fig. 2). To our knowledge, these are the first high-resolution structures of these assemblies determined directly from non-recombinant, native IgM in patient serum, and the first structural estimate of their relative proportions within a single sample, distinct from bulk compositional estimates of CD5L incorporation derived from mass spectrometry of pooled serum^21^. These proportions should be read as illustrative rather than absolute, however, sample preparation and image-processing choices can each skew apparent class distributions. Circulating SC and secretory IgM have previously been associated with liver disease and hepatobiliary injury^31–33^, but one structural dataset per clinical context cannot establish a disease-stage association. Larger cohorts must determine whether IgM composition tracks infection phase, liver injury, viral burden or outcome.

Our emphasis on IgM reflects the assemblies accessible to structural analysis in the severe acute sample, rather than an assumption that IgM predominates over IgG in chronic HBV infection. The IgG-like SVP contacts and two-particle bridges are consistent with a contribution of IgG to particle organization (Supplementary Fig. 2), but provide too few identifiable events to compare network-forming capacity across isotypes. IgG’s smaller size may reduce its detectability by cryo-ET, while enrichment, particle selection and differences in antibody abundance in the recovered fractions may also affect the observed frequencies.

The central advance is the organization that emerges when a polymeric pentameric antibody meets a repetitive, multi-epitope decoy particle. Intraparticle multivalency enhances IgM avidity through simultaneous engagement of HBsAg on the same SVP by multiple Fab arms of a single IgM, stabilizing antibody-particle binding even when individual Fab-HBsAg interactions have low affinity^35,36^. Interparticle bridging extends this principle across particles, linking multiple SVPs through shared IgM contacts into crosslinked networks (Figs. 3 & 4). This avidity-driven crosslinking, rather than higher-affinity individual interactions, likely explains how IgM organizes the repetitive SVP surface into the higher-order architecture described here, extending the classical decoy model beyond passive sequestration (Supplementary Fig. 17). Whether this organization alters virion neutralization or immune-complex clearance remains to be tested. Because enrichment can bias which circulating complexes are preserved, these reconstructed networks should not be treated as an unperturbed map of whole blood.

The observed network containing 10 IgM molecules and 10 SVPs spans approximately 200 × 100 nm (Fig. 4a-b), compared with 22-28 nm for individual SVPs (Supplementary Fig. 15). Ganesan et al. measured ovalbumin-IgG complexes by dynamic light scattering (DLS), reporting volume-weighted mean diameters of approximately 39 nm (range 20-150 nm) at 15-fold antigen excess, 220 nm at threefold excess and 1.23 µm at equivalence^37^. Their distinction between small complexes and antibody-coated bacteria or blood cells places our network in context: crosslinking enlarges SVPs into a nanoscale assembly, without establishing a bacteria-sized “large” immune complex^37^. Turman et al. measured IgG-opsonized HIV-like particles at modal diameters of approximately 134 and 143 nm by nanoparticle tracking analysis and demonstrated FcγRIIb-mediated clearance by liver sinusoidal endothelial cells in humanized mice^38^. Together, these studies place the observed IgM-SVP network on a physical scale relevant to hepatic uptake, but do not establish an IgM clearance pathway. Whether crosslinking promotes removal or tissue deposition requires direct measurements of complex-size distributions, complement deposition and cellular uptake.

SVP-associated IgM adopts distinct Fab-arm conformations with different potential for complement engagement. Planar, stapled and hybrid states coexist (Fig. 4c-d), with the stapled state likely compatible with C1q engagement as described in model systems^11,12^. Independently of whether an IgM engages one SVP or bridges several, a subset of SVP-bound IgM adopts this stapled conformation, which could engage complement and trigger clearance mechanisms directed against the complex. HBsAg-antibody-complement complexes have been implicated in HBV-associated extrahepatic disease, including polyarteritis nodosa and membranous nephropathy^39,40^. However, this stapled conformation was uncommon, and without resolved C1q or an activation assay, we have no evidence that these assemblies activate complement in vivo. More broadly, this study extends structural serology from identifying serum antibodies to resolving how full-length antibodies organize pathogen-derived particles^28,29^. For HBV, the extraordinary SVP burden is therefore not only an antibody sink but also a multivalent platform on which serum IgM assembles higher-order immune complexes.

## Supporting information

Supplementary figures, movie list, and tables

## Acknowledgements

We thank Axel Brilot, Evan Schwartz, and Michelle Mikesh from the University of Texas College of Natural Sciences and award RR160023 of the Cancer Prevention and Research Institute of Texas for support of the EM facility at the University of Texas at Austin. The SBGrid Consortium provided computational support for structural biology software. A portion of this research was supported by NIH grant R24GM154185 and performed at the Pacific Northwest Center for Cryo-EM (PNCC) with assistance from Claudia López. This work was funded by the Cancer Prevention and Research Institute of Texas (RR230050) (ZK), Welch Foundation (F-2244-20250403) (ZK), CNS-catalyst (CAT-26-KE) (ZK), and Lyda Hill Foundation (ZK). The work of AD, SS, SN, SU and RB were funded by the German Research Foundation (DFG) through the TRR179 (project number 272983813; subproject TP23 to SS; subproject TP15 to SN and SU; subproject TP09 to RB).

## Author contributions

CL, SS, and ZK conceived the study. CL, AD, XY, JP, GP, SS, and ZK designed the study. AD enriched the serum samples with HBsAg and IgM and performed characterization, with the supervision of SS. CL prepared cryo-EM grids and performed grid screening. CL and SM collected all the cryo-EM and cryo-ET datasets, with the supervision of ZK. CL and ZK processed the single-particle cryo-EM data, and CL built the PDB models and performed the structural analysis. CL, XY, and ZK processed the cryo-ET data. XY performed subtomogram averaging and geometric-based interaction prediction from tomograms, with the supervision of ZK. GP performed Monte Carlo randomization simulations, with inputs from XY, CL, and ZK. CL and JP designed and performed the ELISA experiments, with the supervision of GG. SN contributed patient serum samples (or reagents). SU and RB contributed in reagents, analysis tools, and instruments. CL and ZK prepared the initial draft. CL, AD, XY, GP, SS, and ZK wrote and revised the manuscript with input from all authors. CL, XY, AD, and GP prepared the figures, tables, and movies. SS and ZK acquired funding and managed the project.

## Materials and Methods

### Ethics statement

Use of human clinical samples was approved by the Ethics Committee of the Medical Faculty of Heidelberg University (approval no. S-153/2024) and conducted in accordance with the Declaration of Helsinki. Written informed consent was obtained from all patients whose samples were collected prospectively.

### Patient serum samples

Serum samples used for cryo-EM and cryo-ET data collection were obtained from two male patients with HBV infection. The patient with severe acute hepatitis B was 83 years old at the time of enrollment on March 21, 2024. He was diagnosed with acute HBV infection or reinfection and died the following day from complications associated with acute hepatitis B. At the time of sampling, he was infected with HBV genotype D1 and had a serum HBsAg concentration of 37,325 IU/mL and an HBV DNA level of 1.09 × 10^8^ IU/mL. Serum aspartate aminotransferase (AST) and alanine aminotransferase (ALT) levels were 395 and 772 U/L, respectively, and liver stiffness measured by FibroScan was 12.6 kPa. The patient with chronic HBV infection was 36 years old at the time of enrollment on September 19, 2024. He was infected with HBV genotype D1 and had a serum HBsAg concentration of 30,120 IU/mL and an HBV DNA level of 212,000 IU/mL. Serum AST and ALT levels were 16 and 35 U/L, respectively, and liver stiffness measured by FibroScan was 6 kPa. Additional demographic and clinical information are provided in Supplementary Table 1.

### Purification of HBV virions and subviral particles

HBV virions and subviral particles were enriched by ultracentrifuging the HBV positive sera into a sucrose cushion. Briefly, 3 mL of 60% (w/w) sucrose was overlaid with 3 mL of 30% (w/w) sucrose, both prepared in TN buffer (20 mM Tris-HCl, 140 mM NaCl, pH 7.4), followed by 18 mL of serum. Samples were centrifuged at 28,000 rpm (107,811 × g) for 15 hr at 4°C. Fractions à 0.5 mL were collected from the bottom and characterized by anti-HBs ELISA and SDS-PAGE followed by Coomassie staining or anti-HBsAg western blotting as described previously^30^.

All chromatography runs were performed at 4°C on an ÄKTA chromatography system (GE Healthcare). Runs were monitored in real-time by measuring UV absorption at wavelengths of 254 and 280 nm. To separate the virion particles from other large proteins found in the serum, 3.5 mL of pooled fraction was subjected to size exclusion chromatography (SEC) on a Superose 6 column (Ø 2.6 cm, length 55 cm).

SEC fractions containing viral particles were subsequently pooled and subjected to heparin affinity chromatography (HAC). In brief, 22 mL of fractions from SEC were applied to 1ml HiTrap Heparin HP column (GE Healthcare). After sample application, the columns were washed with 5 column volumes TN buffer. Column-bound material was eluted by a linear salt gradient over 10 column volumes ranging from 140 to 2,140 mM NaCl in 20 mM TrisCl (pH 7.4). Elution fractions à 0.5 mL were immediately collected and processed further.

The concentrated peak fractions obtained from HAC were subjected to equilibrium density gradient ultracentrifugation. 1.5 mL of HAC eluate was layered on top of a preformed discontinuous sucrose step gradient (Sucrose w/w in TN buffer: 1 mL 60%, 0.5 mL each of 45%, 35%, 25%, 15%) and centrifuged at 55,000 rpm (406,849 × g) for 15 hr at 4°C. Fractions were collected from the bottom in volumes à 6 drops (∼150 μL).

### Western blots

Fractions were mixed with 6 × Laemmli buffer (97.5 mM TrisCl (pH 6.8), 30% glycerol, 3%SDS, 7.5% β-mercaptoethanol, 0.06% (w/v) bromophenol blue) and boiled at 65°C for 30 min. Fractions were loaded in 12% SDS-PAGE gel and transferred to Immobilon-FL membrane (Merck Millipore, IPFL00010) using Trans-Blot Turbo transfer system (Bio-Rad, USA). The membranes were blocked with 5% BSA for 1 h at room temperature, followed by incubation with human monoclonal anti-HBs antibody (HBD87, Humabs Biomed) overnight at 4°C. The following day, the membranes were washed with PBST and incubated with polyclonal goat anti-human IgG antibody conjugated to IRDye 680RD (LICOR, 926-68078) for 1 hr at room temperature. The membranes were visualized using Odyssey M Imaging System (LICOR). Alternatively, membranes were incubated with polyclonal goat anti-human IgG antibody conjugated to HRP (abcam, ab97165). Subsequently, signals were visualized with Clarity Western ECL substrate (Bio-Rad, 1705061) or Amersham ECL Prime Western Blot Detection Reagent (Cytiva, RPN2232) on ChemiDoc (BioRad).

### ELISA analysis of serum IgM, CD5L-associated IgM, and SC-associated IgM

Serum total IgM, CD5L-associated IgM, and SC-associated IgM were evaluated using ELISA. ELISA assays were done in a similar way with our previous work^41^. Serum samples used for ELISA were obtained from BioCollections Worldwide, Inc. Sera from healthy donors were confirmed to be non-reactive for HIV, HBV, and hepatitis C virus (HCV). Sera from HBV-positive donors were reactive for HBsAg and non-reactive for both anti-HIV and anti-HCV. The HBV-positive serum samples were collected in serum separator tubes from six individual donors. Two donors were male residents of the United States, whereas sex was not specified for the remaining four donors, who were residents of Guinea-Bissau. HBV DNA levels ranged from 2.80 × 10^8^ to 7.58 × 10^8^ IU/mL and were quantified using the Aptima HBV Quantitative Assay. HBsAg reactivity was determined using the Genetic Systems HBsAg EIA 3.0 assay. Additional sample characteristics, including collection dates, test dates, and individual HBV viral loads, are provided in Supplementary Table 2.

High-binding 96-well plates were coated separately with 50 μL per well of anti-human CD5L (Thermo Fisher Scientific, PA5-20243; 5 μg/mL), anti-human SC (Thermo Fisher Scientific, MAHU-SC-MAB; 5 μg/mL), or anti-human IgM (Thermo Fisher Scientific, MA5-37963; 4 μg/mL), each diluted in PBS. Plates were sealed and incubated overnight at 4°C. The following day, coating solution was removed and plates were washed three times with PBST (PBS containing 0.05% Tween-20). Nonspecific binding was blocked with PBSTM (PBST containing 5% nonfat dry milk; 200 μL/well) for 3 h at 37°C, followed by three washes with PBST. Serum samples were serially diluted twofold in PBSTM, from 1:2 to 1:2,048 for anti-CD5L and anti-SC assays, and from 1:4 to 1:4,096 for the anti-IgM assay. PBSTM alone was used as the negative control. 50 μL of each dilution were incubated on the coated plates for 1 h at 37°C. After three washes with PBST, captured IgM was detected using horseradish peroxidase (HRP)-conjugated anti-IgM antibody (Southern Biotech, 2020-05; 1:5,000 in PBSTM; 50 μL/well) for 1 h at 37°C. Plates were washed four times with PBST, followed by 50 μL of tetramethylbenzidine (TMB) substrate (Thermo Fisher Scientific, 34028) added per well. The reaction was quenched with 50 μL of 2 M sulfuric acid, and absorbance was measured at 450 nm using a microplate reader. Data are reported as raw absorbance values (OD450).

### Cryo-EM and cryo-ET grid preparation

Sample vitrification was performed using a Leica EM GP2 plunge freezer operated at 20 °C and 90% humidity. A total of 3 μL of sample was applied to holey carbon grids (C-flat 300 mesh 2/2 grid) that had been glow-discharged for 30 s with Leica ACE600. The grids were back-side blotted for 3 s using standard filter paper and then plunge-frozen in liquid ethane.

### Cryo-ET data collection

The severe acute sample was used for cryo-ET data collection using two SEC fractions with high and low particle concentrations. For the high-concentration fraction, frozen grids were screened on a Talos Glacios, and selected grids were shipped to the Pacific Northwest Cryo-EM center (PNCC) for data collection. Tilt series were acquired on a Titan Krios transmission electron microscope (Thermo Fisher Scientific) operating at 300 kV, equipped with a 10-eV slit width energy filter and a Falcon 4i direct electron detector. Data were collected using Tomo5 (Thermo Fisher Scientific) at a nominal magnification of 81,000×, corresponding to a pixel size of 1.512 Å. A total of 269 tilt series were acquired using a dose-symmetric tilt scheme from −60° to +60° in 3° increments, with a total accumulated dose of ∼120 e^−^/Å^2^ and a defocus range of −2 to −6 μm^42^. Data collection details are summarized in Supplementary Table 3. For the low-concentration fraction, tilt series were acquired on a Talos Glacios at a nominal magnification of 73,000×, corresponding to a pixel size of 2.01 Å. Data were collected using SerialEM^43^ with a dose-symmetric tilt scheme from −60° to +60° in 3° increments, a total accumulated electron exposure of ∼120 e^−^/Å^2^, and a defocus range of −4 to −6 μm. Detailed data acquisition parameters are summarized in Supplementary Table 3.

### Cryo-ET data processing

For subtomogram averaging of the high-concentration fraction, tilt series data processing was carried out using RELION 5^44^. Raw EER frames and mdoc files were imported, followed by motion correction using MotionCor2^45^ and CTF estimation using CTFFIND4^46^. Tilt series alignment was then performed using IMOD^47^ patch tracking after low-quality tilt images were manually excluded. Tomograms were subsequently reconstructed in RELION 5 at 6× binning (9.072 Å/pixel) using the resulting alignments.

To improve tomogram visualization for Fab arm picking, 8 tilt series from the high-concentration fraction and 22 tilt series from the low-concentration fraction were processed using WarpTools^48^ and MissAlignment^49^. Raw EER frames and mdoc files were imported into WarpTools, where motion correction and initial CTF estimation were performed. An initial coarse tilt series alignment was generated by patch tracking in IMOD, followed by tilt series CTF estimation in WarpTools. MissAlignment was then applied to refine the tilt series alignment, after which tilt series CTF estimation was repeated using the updated alignment parameters. Tomograms were subsequently reconstructed in WarpTools at 6× binning (9.072 Å/pixel) using the refined alignments.

To further improve tomogram quality for visualization and particle picking, the tomograms were processed with IsoNet2, a deep learning-based tool for missing-wedge compensation and denoising^50^. For IsoNet2 model training, 10 selected tomograms were used for each dataset. Then, the denoising model was first trained and applied to the selected tomograms, followed by training of an initial refinement model to generate higher-quality tomograms for mask generation. Masks were then generated from these tomograms and used to train a final refinement model. The corresponding final model was subsequently applied to all tomograms within each dataset.

The final tomograms processed by MissAlignment and IsoNet2 show reduced background noise and improved the definition of particle densities in the reconstructed tomograms, facilitating subsequent particle identification, segmentation, and three-dimensional visualization.

### Particle picking using crYOLO

To facilitate manual particle picking for crYOLO model training, IsoNet2-processed tomograms from the RELION 5-reconstructed high-concentration dataset were used. A total of 1,012 IgM particles from 10 tomograms and 1,051 SVP particles from 4 tomograms were manually picked using the Napari GUI to generate initial training datasets for crYOLO^51^, a deep learning-based particle picking tool. Using the manually picked particle lists as input, an initial model was trained and applied back to the selected tomograms to enable more comprehensive particle recognition. The resulting particle picks were then supplemented by manually picking remaining missed particles. A final model was then trained and applied to all tomograms in the dataset. In total, 65,504 SVP and 95,302 IgM particles were picked for later analysis (Supplementary Figs 3, 15).

### Subtomogram averaging

Subsequent processing was performed in RELION 5^44^. To generate an initial reference and mask for 3D refinement, 217 IgM particles were manually picked with approximate positions and Euler angles using the UCSF ChimeraX ArtiaX plugin^52^, followed by particle extraction and subtomogram reconstruction. The resulting reconstruction was used as an initial reference for subsequent processing of the IgM particles identified by crYOLO.

All IgM particles picked by crYOLO were then extracted at 4× binning (6.048 Å/pixel, box size 128 pixels). Standard 3D auto-refinement and 3D classification were then performed to remove falsely picked particles. A total of 28,035 particles remained and were re-extracted at 2× binning (3.024 Å/pixel, box size 168 pixels) to achieve higher resolution for IgM assembly identification. After several additional rounds of 3D refinement and classification using updated references and masks, 18,368 particles remained and were classified into three IgM assemblies, IgM-J-CD5L, IgM-J, and IgM-J-SC. The three assemblies were resolved to 7.6 Å, 11.4 Å, and 8.8 Å, respectively. The refined particle positions and Euler angles were subsequently used to map individual IgM particles back into the tomograms for interaction analysis. (Supplementary Fig. 3).

A similar processing strategy was applied to SVP. All 65,504 SVP particles picked by crYOLO were extracted at 4x binning (6.048 Å/pixel, box size 96 pixels). An octahedral symmetry reconstruction generated was used as the initial reference for 3D classification. After 3D auto-refinement and 3D classification to remove falsely picked particles, 61,963 SVP particles remained for further classification, resulting in 7 SVP classes distinguished particle diameter, ranging from 22 to 28 nm in diameter. (Supplementary Fig. 15).

### Segmentation

For segmentation of tomograms, different particle-localization strategies were used depending on particle concentration. For high-concentration tomograms, the subtomogram averaging refined coordinates of SVPs were imported into ArtiaX^52^ in UCSF ChimeraX. See Supplementary Fig. 15 for subtomogram averaging workflow. Because the SVPs are heterogeneous, we manually segmented the full SVPs identified above; this is achieved by placing the markers within each positioned particle volume. For low-concentration tomograms, individual IgM molecules and SVPs were readily identified from their tomographic densities, and markers were placed directly within each particle density. In both cases, a separate group of markers was assigned to each IgM/SVP and used to segment individual particle densities from the tomograms using the Color Zone, followed by the Split Map in ChimeraX.

### Visualization of IgM in tomograms

For visualization of IgM in high-concentration tomograms, the refined coordinates and orientations of IgM Fc cores obtained from subtomogram averaging were imported into ArtiaX in UCSF ChimeraX. The corresponding subtomogram averages were placed back into the tomograms according to their refined coordinates and orientations. The appropriate class average was placed at each particle position according to its assignment as IgM-J-CD5L, IgM-J-SC, or unbound IgM-J, allowing the compositional state of individual IgM molecules to be visualized.

Placement of the Fc-core averages provided spatial references for localizing the Fab arms extending from each IgM molecule. For visualization of the IgM Fab arms, a structural model containing the Cμ1, Cμ2, VH, VL, and CL domains, extracted from a previously reported human IgM structure (PDB 8ADZ^19^), was used as the Fab reference model. Given the inherent flexibility of the Fab arms, this model was used to approximate their overall positions and orientations rather than to represent their precise conformations in the tomograms. Fab arms were manually identified based on their densities in the tomograms, with the hinge positions defined by the IgM Fc core providing additional guidance. The Fab reference model was then manually positioned and oriented in ArtiaX to match the corresponding tomographic densities. Because IgM has five Fab arms, and sometimes we could not reliably detect all of them, so the missing Fab arm positions were arbitrarily placed while not colliding the exisiting IgM or SVP densities. This is for visualization purpose, and they are labeled in gray in the figures to separate them from the reliable Fab arm densities (salmon).

### Direct observation of IgM-SVP interactions in tomograms

IgM-SVP interactions were directly assessed in the tomograms using the IgM and SVP visualizations described above. Positioned IgM Fc-core averages provided the locations and orientations of individual IgM molecules and guided localization of their Fab arms, while segmented SVP densities defined the positions and surfaces of individual SVPs. The Fab arms were manually traced based on their tomographic densities using the UCSF ChimeraX ArtiaX plugin. IgM-SVP interactions were then directly identified based on the spatial relationship between the traced Fab arms and the surfaces of neighboring SVPs.

### Calculation of Fab arm angles

To characterize the spatial relationship between the IgM core and Fab arms, the in-plane angle, out-of-plane angle (both measured relative to the IgM’s own equatorial plane), and spin angle (the Fab arm’s own rotation about its axis) of the 180 Fab arms were recorded. Two outliers, with an in-plane angle of approximately 72° and an out-of-plane angle of approximately 130°, were identified and excluded, leaving 178 confirmed measurements. Based on the distribution of these measurements, the in-plane angle range was ±60°, the out-of-plane angle range was ±110°, and the spin angle range was ±180° (Supplementary Fig. 14). The hinge position (the junction between Cμ2 and Cμ3) were identified in ChimeraX for each IgM assembly and defined as Fab anchor sites. Based on these anchor positions, the reachable space of each anchor site was modeled as a solid elliptical cone with a depth equal to the Fab arm length (12.5 nm) (Supplementary Fig. 16a).

### Geometry-based prediction of IgM-SVP interactions from tomograms

For prediction of IgM-SVP interactions, quality control was first performed to select tomograms and particles suitable for quantitative analysis. All 253 tomograms were manually reviewed, and those showing an implausible ratio of IgM to SVP particle counts, indicating likely artifacts in particle picking or segmentation, were excluded, leaving 133 tomograms for downstream analysis. Within each tomogram, a particle was counted toward the interaction statistics only if its distance from the XY boundary was large enough to ensure that any interaction partner it might have would not fall outside the tomogram; a particle closer to this boundary could still be tested as the interaction partner of a qualifying particle on the opposite side, but was not itself counted, since a genuine partner located just outside the tomogram could not be captured. This boundary margin was applied only in the XY plane; Z-boundaries were not excluded, since the ice layer imaged here is thin.

Interactions between IgM and SVPs were determined using the following probabilistic method. A two-dimensional kernel density estimate (KDE) fit to the in-plane (±60°) and out-of-plane (±110°) angles of the 178 confirmed Fab hinge angle measurements described above (Supplementary Fig. 14). Because each anchor site corresponds to a pair of Fab arms, whereas the reachable space was modeled for an individual Fab arm, an angular offset of 18.5° between the central axis of the Fab pair and the axis of an individual Fab was added to the observed half-ranges, resulting in final angular limits of ±78.5° in-plane and ±128.5° out-of-plane. The resulting set of angles was discretized onto a fixed 2° angular grid, restricted to the cone’s elliptical angular range (in-plane half-angle 78.5°, out-of-plane half-angle 128.5°), and renormalized so that its weights summed to 1 over the retained directions.

For each Fab anchor site, a reach probability was computed as the sum of the grid weights over all candidate arm directions whose full-length (12.5 nm) arm segment came within the SVP’s radius of the SVP center. An IgM particle was scored as interacting with an SVP if at least one of its five anchor sites had a reach probability exceeding a threshold of 0.05. This 0.05 threshold was calibrated empirically: IgM-SVP interactions were first identified manually in one representative tomogram, and the automated pipeline was then run across a range of threshold values, comparing how closely the resulting interaction counts tracked the manually identified ones, with 0.05 giving the closest agreement. See more details in Supplementary Fig. 16a.

### Monte Carlo simulation of IgM-SVP spatial interactions

IgM-SVP spatial interactions were modeled using a custom Python 3 program and deposited at github with the link here: https://github.com/Ke-Lab-UT-Austin/IgM-SVP_Simulations. Specifically, each IgM particle was represented as a rigid pentameric hub with five radiating arms positioned at 0°, 60°, 120°, 180°, and 240°, and each SVP was represented as a sphere decorated with surface spike glycoproteins. Particle dimensions were parameterized based on experimentally estimated values, including an IgM hub radius of 8.6 nm, arm length of 7.5 nm, Fab-tip radius of 2.5 nm, SVP radius of 9.5 nm, and SVP spike diameter of 2.5 nm (Supplementary Fig. 10a).

An IgM-SVP pair was scored as interacting when the center-to-center distance was less than or equal to the sum of their maximal physical reaches, corresponding to a contact distance of 33.1 nm under the default parameters. No additional contact tolerance was applied. This criterion represents a static geometric contact test and does not model binding kinetics or experimentally observed Fab angular distributions. The formula used to compute binding is shown below:

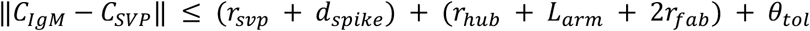

Each IgM was permitted to interact with up to 10 distinct SVPs, whereas no limit was imposed on the number of IgM molecules that could interact with a single SVP. Nearest-neighbor searches were accelerated using SciPy’s KDTree.

To generate a spatial null model, IgM and SVP centers were randomly placed within the simulation volume by uniform rejection sampling subject to steric-exclusion constraints that prevented nonphysical particle overlap. IgM and SVP populations were placed in an interleaved manner, and each IgM was assigned a random azimuthal rotation about the z-axis. Placement was attempted up to 10,000 times per particle, with unconstrained placement used as a fallback when no valid position was found. For comparison with experimental cryo-ET data, IgM and SVP particle coordinates were manually picked in ArtiaX^52^ from the low-concentration tomograms and exported as RELION-format STAR files. IgM and SVP coordinates from each tomogram were converted to nanometers and jointly origin-shifted to match the coordinate convention of the simulated configurations. Real IgM particles were assigned random azimuthal orientations because these orientations were not resolved by particle picking alone. The same geometric interaction criterion was then applied to both experimental and simulated particle configurations.

For each tomogram dimensions and particle density, 100 independent randomized configurations were generated and analyzed. Distributions of the number of interactions per IgM and per SVP were grouped into fixed bins (0, 1, 2, 3, 4, and ≥5), and the mean and standard deviation of the percentage of particles in each bin were calculated across simulations. Simulated distributions were compared with the corresponding experimental tomogram using z-scores and associated P values, together with a chi-square test for homogeneity. Simulations and statistical analyses were implemented in Python 3 using NumPy, SciPy, Pandas, and PyVista, with NumPy’s PCG64 generator used for random sampling without a fixed seed by default. See Supplementary Fig. 10 for more information.

### Cryo-EM data collection

One dataset was collected from the chronic sample, and two datasets were collected from different SEC fractions of the severe acute sample. Patient information is described above and summarized in Supplementary Table 1. Frozen grids were screened on a Talos Glacios. The optimal girds were shipped to the PNCC for data collection. Movie stacks were acquired using a Titan Krios transmission electron microscope (Thermo Fisher Scientific) operating at 300 kV, equipped with a 10-eV slit width energy filter and a Falcon 4i direct electron detector. Images were recorded using EPU (Thermo Fisher Scientific) at a nominal magnification of 130,000x in counting mode, corresponding to a pixel size of 0.93 Å. The total electron exposure was 50 e^−^/Å^2^. The defocus range was set from −0.7 to −2.4 μm. Data collection details are summarized in Supplementary Table 4.

### Cryo-EM data processing

Movies were imported into cryoSPARC^53^ for processing. Motion correction and patch CTF estimation were performed in cryoSPARC 4.7.1. Particles were automatically picked using the blob-picking, followed by particle extraction and several rounds of 2D classification to remove contamination and poorly aligned particles. The blob-picked dataset was dominated by SVP particles, with a smaller population of IgM particles identified by 2D classification. An initial reconstruction using ∼19,000 IgM particles yielded a map at approximately 8 Å resolution. Because top-view IgM particles are thin and prone to misclassification during 2D classification, the ∼19,000 initially identified IgM particles were subjected to C50 symmetry expansion to generate multiple in-plane orientations, yielding ∼950,000 expanded particles. These particles were included during 2D classification of the entire blob-picked dataset (∼5 million particles), which enabled recovery of additional IgM particles that had initially been overlooked. Duplicate particles were removed in the final stage of 2D classification. The recovered particles produced 2D class averages consistent with the pentameric IgM architecture (Supplementary Fig. 5).

All IgM particles were first reconstructed together, yielding a map corresponding to the IgM-J-CD5L complex. Subsequent 3D classification revealed additional structural classes. To further separate heterogeneous particles, several structural models were used as references, including IgM-J-CD5L, IgM-J, and IgM-J-SC. The classification results indicated that the particle population consisted primarily of IgM-J-CD5L, IgM-J, and IgM-J-SC complexes. These three models were then used as starting references for heterogeneous refinement of the entire IgM particles in cryoSPARC. Several additional rounds of 2D classification and 3D classification were performed to further remove contamination and improve particle homogeneity. The final particle subsets were subjected to non-uniform refinement. Reference-based motion correction was then applied to improve particle alignment and map resolution. Finally, spIsoNet^50^ was used for refinement of the final reconstructions (Supplementary Figs. 4-5). DeepEMhancer^54^ was used for whole-map visualization, whereas EMReady2^55^ was used to enhance side-chain features for visualization and interpretation. Additional structural models, including IgM hexamer^56^, FcμR-bound IgM^57,58^, and C1q-bound IgM^11^, were also tested as references during heterogeneous refinement.

### Model building and refinement

Atomic models were built using ISOLDE^59^ in UCSF ChimeraX. Previously reported structures of IgM-J-CD5L (PDB: 8R83)^20^ and IgM-J-SC (PDB: 6KXS)^16^ were rigid-body docked into the cryo-EM density map and used as starting models. Due to the relatively poor local resolution of the Cμ3 regions, individual Cμ3 domains were separated from the starting models and independently rigid-body fitted into the corresponding densities, whereas the Cμ4 domains and tailpieces were subjected to molecular-dynamics flexible fitting in ISOLDE^59^. Regions with unambiguous density were rebuilt where necessary, and glycans were modeled into well-resolved glycosylation sites (Supplementary Fig. 4d). The resulting models were subjected to real-space refinement in Phenix ^60^. Model quality and map-model agreement were evaluated using MolProbity scores, Ramachandran statistics, and map-model FSC. Statistics for model refinement and geometry are summarized in Supplementary Table 5.

### Structural comparison of Fcμ5B in IgM-J and IgM-J-SC

To quantify the observed conformational difference in Fcμ5B between the IgM-J and IgM-J-SC structures, the structures were first aligned using the J chain and Fcμ5A as a common reference. Visual comparison of the aligned structures revealed a change in the orientation of the Fcμ5B region encompassing residues Asn395-Pro401. For each structure, a vector was therefore defined from the Cα atom of Asn395 to the Cα atom of Pro401 in Fcμ5B to represent the orientation of this region. The angle between the Asn395-Pro401 vectors from the IgM-J and IgM-J-SC structures was calculated from their normalized dot product to quantify the change in Fcμ5B orientation. Structural alignment and vector-angle calculations were performed in PyMOL (The PyMOL Molecular Graphics System, Version 3.0.3, Schrödinger, LLC) using custom scripts. In addition, the separation between Fcμ5A and Fcμ5B within each structure was measured in UCSF ChimeraX as the Cα-Cα distance between the corresponding Pro402 residues.

## Data availability

The high resolution cryo-EM structures are deposited at Electron Microscopy Data Bank (EMDB) under accession codes EMD-xxxxx (IgM-J-CD5L, cryo-EM), EMD-xxxxx (IgM-J, cryo-EM), EMD-xxxxx (IgM-J-SC, cryo-EM). The associated atomic models in the Protein Data Bank (PDB) under accession codes PDB-xxxx (IgM-J-CD5L), PDB-xxxx (IgM-J), PDB-xxxx (IgM-J-SC). The cryo-ET maps are deposited at Electron Microscopy Data Bank (EMDB) under accession codes EMD-xxxxx (IgM-J-CD5L, cryo-ET), EMD-xxxxx (IgM-J, cryo-ET), EMD-xxxxx (IgM-J-SC, cryo-ET).

