## Supplementary figures, movie list, and tables for "Serum IgM crosslinks hepatitis B subviral particles into supramolecular immune complexes"

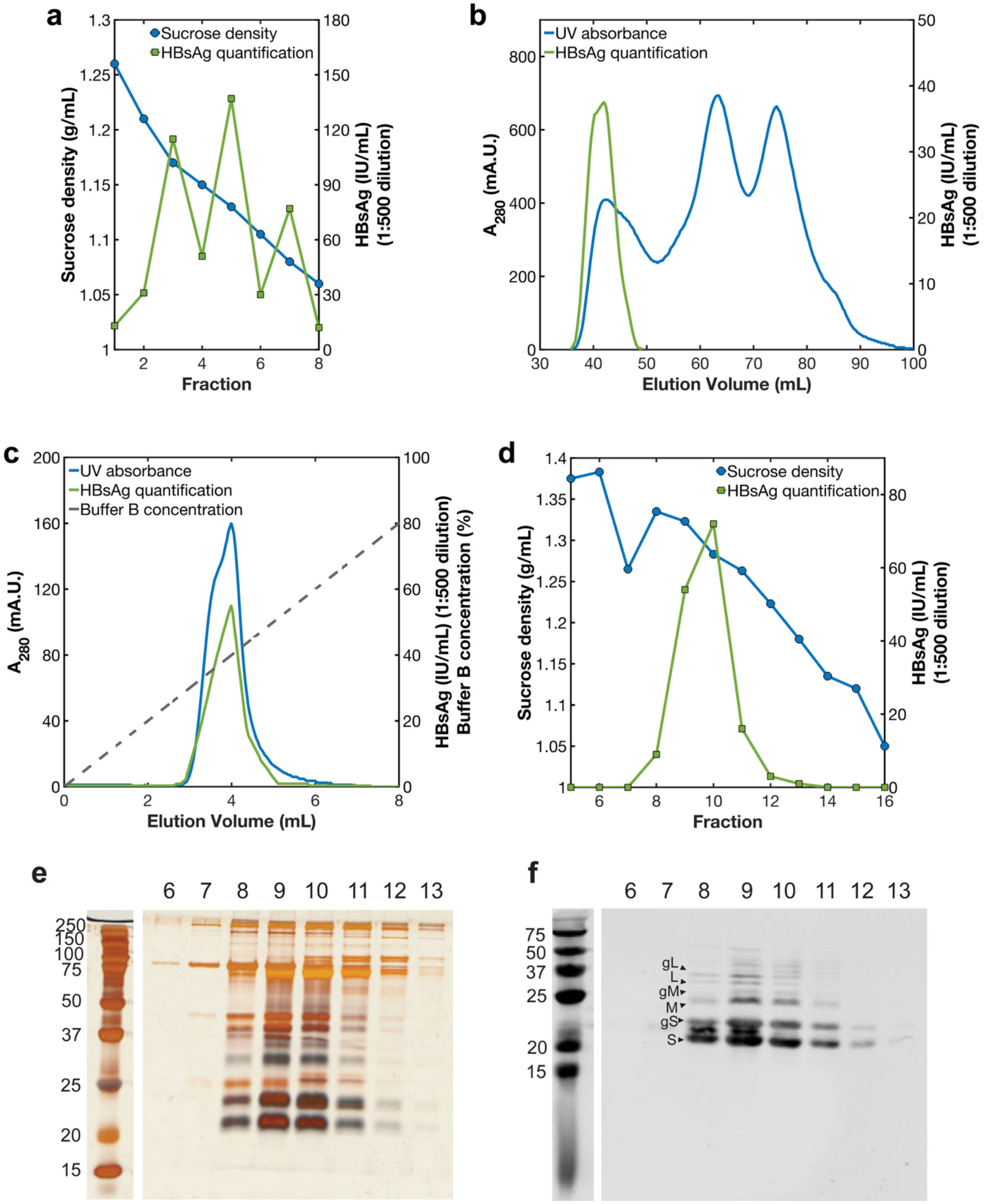

**Supplementary Fig. 1. Purification of HBV from patient serum. a-d.** HBV-positive serum was sequentially processed by **a.** sucrose cushion, **b.** size-exclusion chromatography, **c.** heparin affinity chromatography, and **d.** sucrose gradient centrifugation to purify and enrich HBV viruses and subviral particles. HBsAg levels were measured at each purification step using ELISA. **e.** Silver-stained SDS-PAGE gel showing protein profiles of fractions collected from the sucrose gradient. **f.** Western blot analysis of the same fractions shown in panel E, identifying glycosylated and non-glycosylated forms of HBsAg: glycosylated large (gL), large (L), glycosylated medium (gM), medium (M), glycosylated small (gS), small (S) proteins.

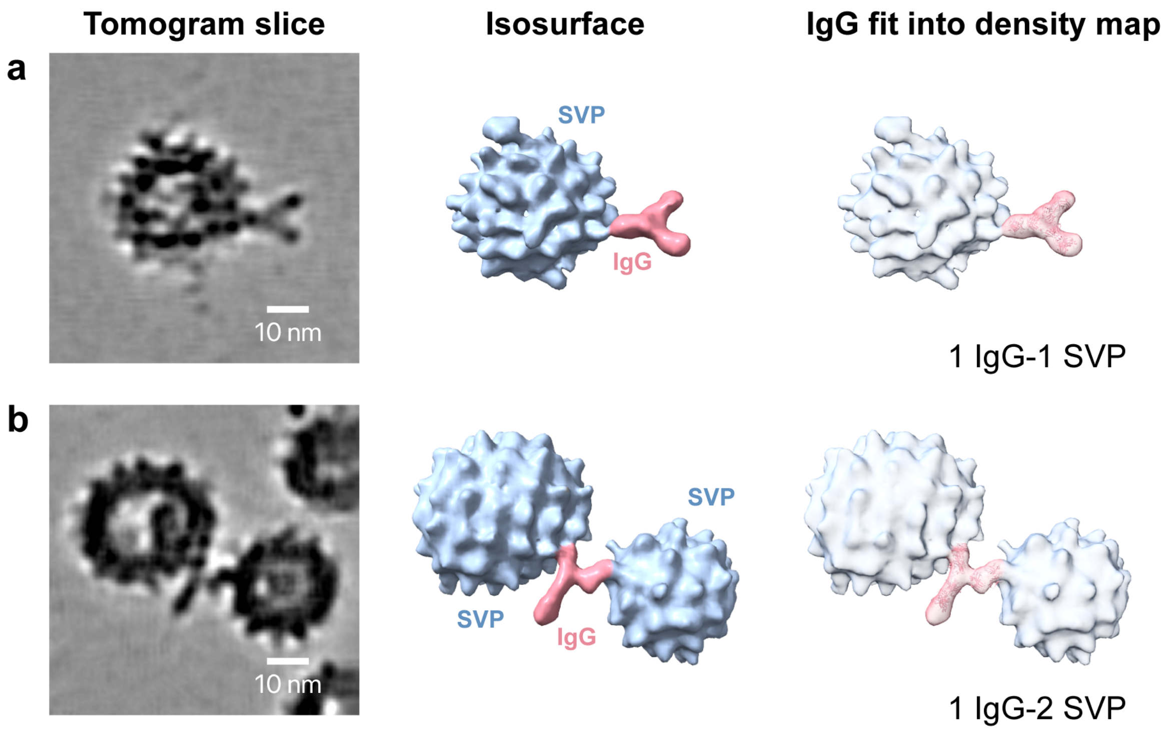


**Supplementary Fig. 2. Cryo-ET reveals IgG-SVP interactions. a-b**, Representative IgG-SVP assemblies illustrating different patterns of engagement, in which individual IgG contact one (**a**) to two (**b**) SVPs. For each assembly, the tomographic slice, corresponding isosurface rendering, and IgG fitted isosurface are shown from left to right. Segmentation colors: SVPs, light blue; IgG, salmon.

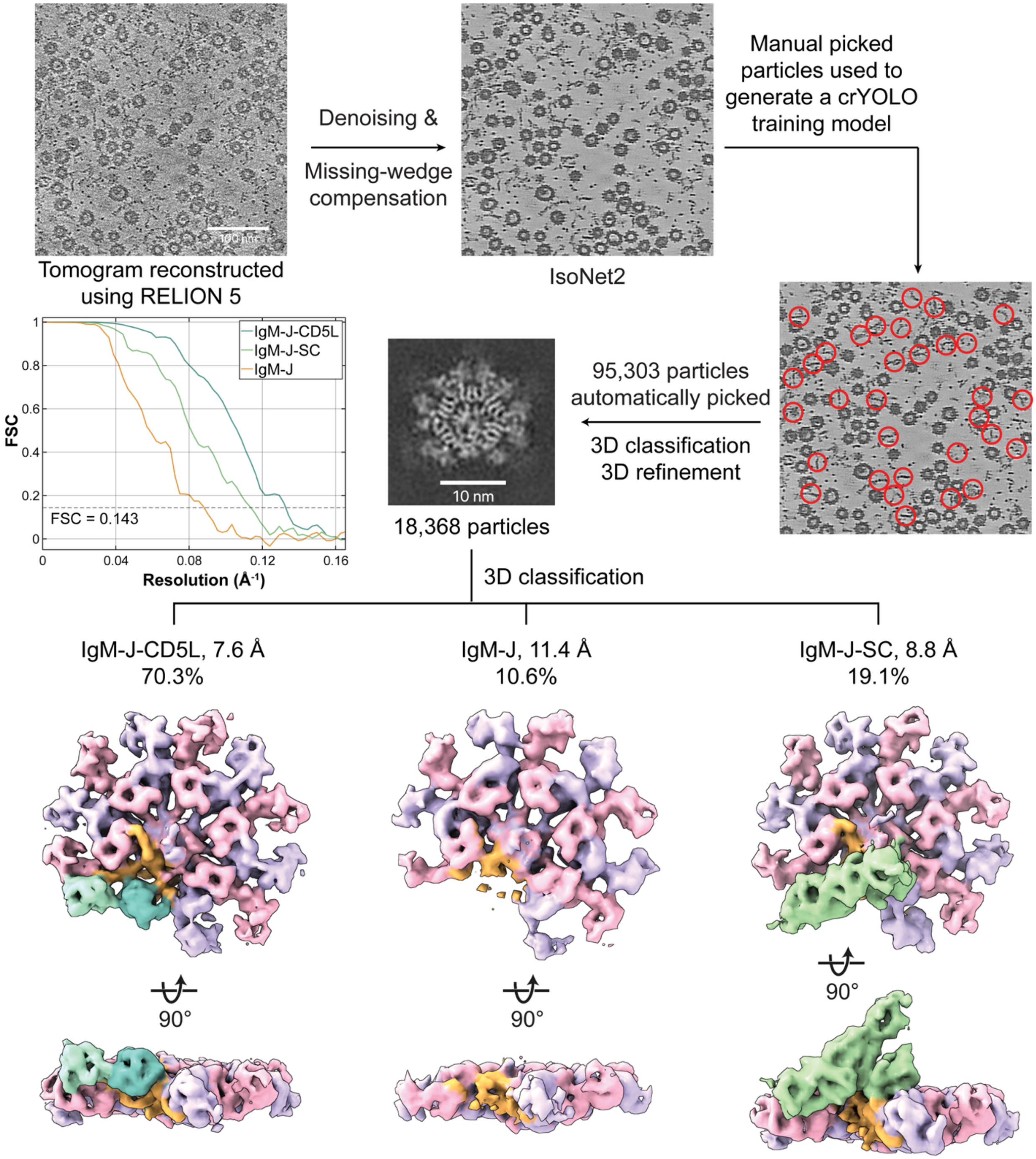


**Supplementary Fig. 3. Cryo-ET data processing workflow for the serum IgM assemblies.** Representative tomographic slice from a tomogram reconstructed using RELION 5. IsoNet2 was applied for denoising and missing-wedge compensation. Manually picked 1,012 IgM particles from 10 tomograms were used to generate a training model for crYOLO, followed by automated picking of 95,303 particles. After 3D classification and refinement, 18,368 IgM particles were selected and subjected to further 3D classification, yielding three distinct IgM assemblies: IgM-J-CD5L, IgM-J, and IgM-J-SC. The resulting subtomogram averages were resolved to 7.6 Å, 11.4 Å, and 8.8 Å, respectively, as determined by the FSC = 0.143 criterion.

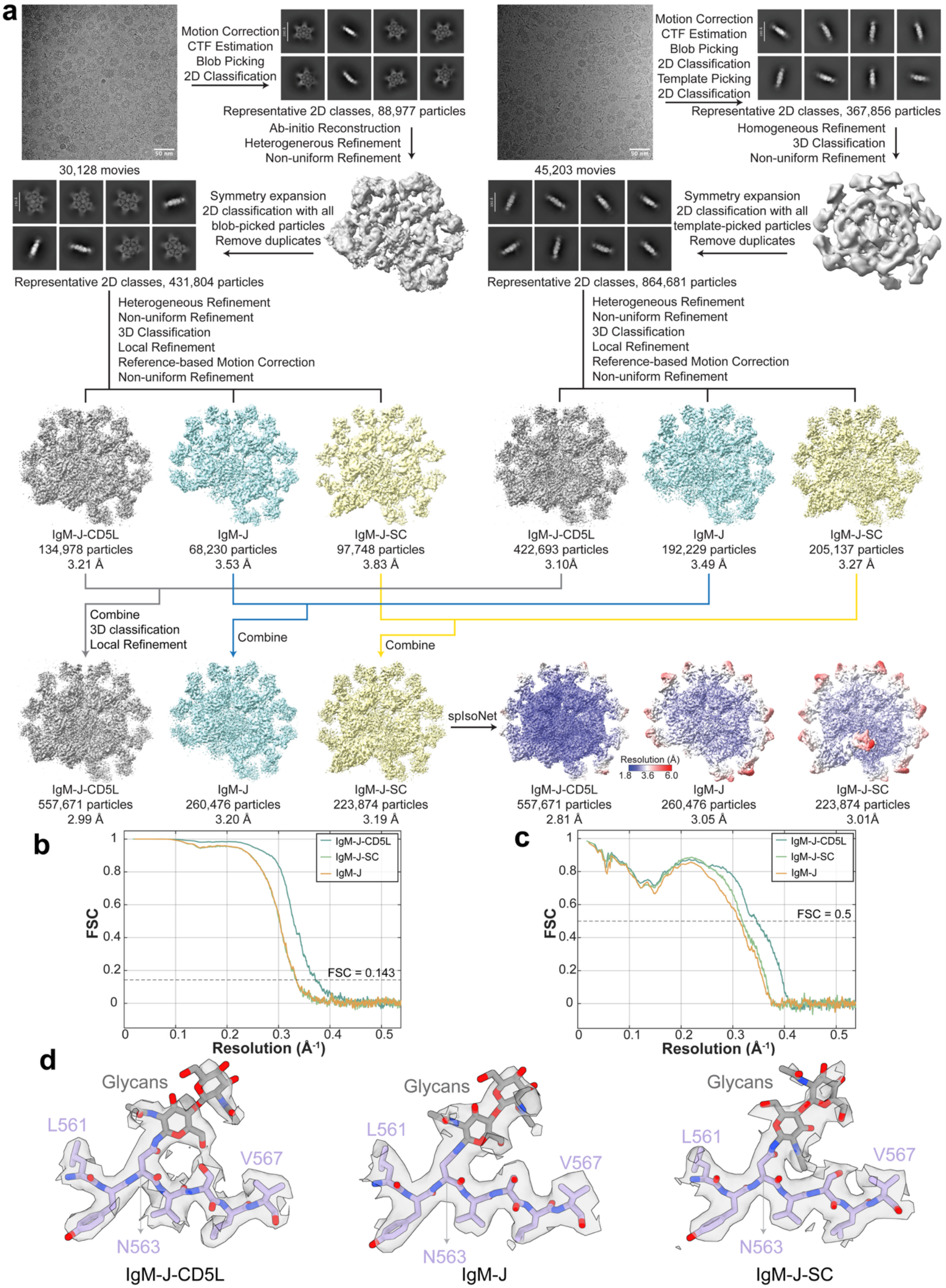
a
Motion Correction
CTF Estimation
Blob Picking
2D Classification
Representative 2D classes, 88,977 particles
30,128 movies
Ab-initio Reconstruction
Heterogenerous Refinement
Non-uniform Refinement
Symmetry expansion
2D classification with all
blob-picked particles
Remove duplicates
Representative 2D classes, 431,804 particles
Motion Correction
CTF Estimation
Blob Picking
2D Classification
Template Picking
2D Classification
Representative 2D classes, 367,856 particles
45,203 movies
Homogeneous Refinement
3D Classification
Non-uniform Refinement
Symmetry expansion
2D classification with all
template-picked particles
Remove duplicates
Representative 2D classes, 864,681 particles
Heterogeneous Refinement
Non-uniform Refinement
3D Classification
Local Refinement
Reference-based Motion Correction
Non-uniform Refinement
IgM-J-CD5L
134,978 particles
3.21 Å
IgM-J
68,230 particles
3.53 Å
IgM-J-SC
97,748 particles
3.83 Å
IgM-J-CD5L
422,693 particles
3.10Å
IgM-J
192,229 particles
3.49 Å
IgM-J-SC
205,137 particles
3.27 Å
Combine
3D classification
Local Refinement
Combine
Combine
IgM-J-CD5L
557,671 particles
2.99 Å
IgM-J
260,476 particles
3.20 Å
IgM-J-SC
223,874 particles
3.19 Å
spIsoNet
Resolution (Å)
1.8 3.6 6.0
IgM-J-CD5L
557,671 particles
2.81 Å
IgM-J
260,476 particles
3.05 Å
IgM-J-SC
223,874 particles
3.01Å
b
IgM-J-CD5L
IgM-J-SC
IgM-J
FSC = 0.143
FSC
Resolution (Å-1)
c
IgM-J-CD5L
IgM-J-SC
IgM-J
FSC = 0.5
FSC
Resolution (Å-1)
d
Glycans
L561
V567
N563
IgM-J-CD5L
Glycans
L561
V567
N563
IgM-J
Glycans
L561
V567
N563
IgM-J-SC

**Supplementary Fig. 4. Cryo-EM data processing workflow for the serum IgM assemblies.** **a,** Two cryo-EM datasets from the same patient serum containing 30,128 and 45,203 movies were collected and then processed separately through motion correction, CTF estimation, particle picking, 2D classification, ab initio reconstruction, heterogeneous refinement, non-uniform refinement, 3D classification, local refinement, and reference-based motion correction. Representative micrographs, 2D class averages, and intermediate reconstructions are shown for each dataset. Classification separated three major IgM assemblies: IgM-J-CD5L, IgM-J, and IgM-J-SC. For the 30,128-movie dataset, refinements yielded IgM-J-CD5L at 3.21 Å from 134,978 particles, IgM-J at 3.53 Å from 68,230 particles, and IgM-J-SC at 3.81 Å from 97,743 particles. For the 45,203-movie dataset, refinements yielded IgM-J-CD5L at 3.10 Å from 422,693 particles, IgM-J at 3.49 Å from 192,229 particles, and IgM-J-SC at 3.27 Å from 205,137 particles. Particles corresponding to the same structural class were then combined and further refined, producing final maps of IgM-J-CD5L, IgM-J, and IgM-J-SC at 2.99 Å, 3.20 Å, and 3.19 Å, respectively. Application of spIsoNet further improved map quality and local density features, yielding final enhanced maps at 2.81 Å, 3.05 Å, and 3.01 Å, respectively. **b,** Fourier shell correlation curves of the half map 1 versus the half map 2 for IgM-J-CD5L, IgM-J, and IgM-J-SC assemblies after processing using spIsoNet, with resolution estimated at the FSC = 0.143 criterion indicated by the dashed line. **c,** Fourier shell correlation curves of refined model versus the full map for the three complexes, with resolution estimated at the FSC = 0.5 criterion indicated by the dashed line. **d,** Cryo-EM density maps of the β-sheet region spanning L561-V567 at the IgM tailpiece and the glycosylation site at N563, with the density of the associated glycans clearly visible.

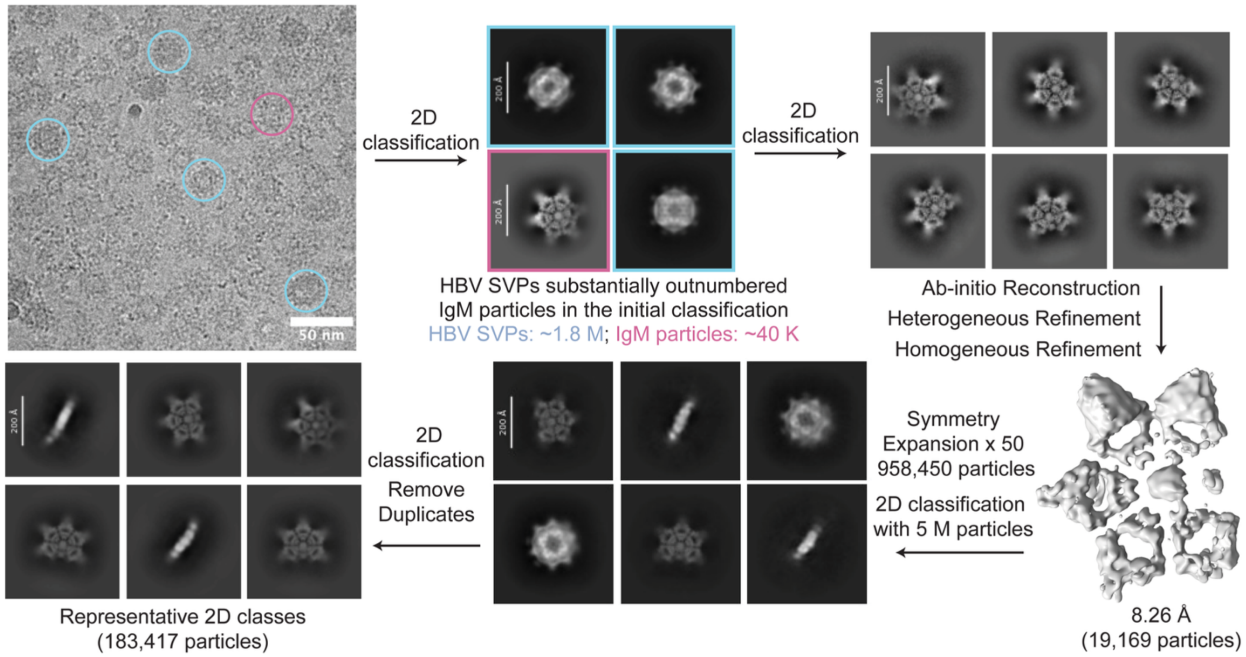


**Supplementary Fig. 5. Symmetry expansion-assisted particle fishing workflow for IgM reconstruction.** Representative cryo-EM micrograph showing abundant HBV SVPs and relatively sparse IgM particles. In the initial 2D classification, HBV SVPs substantially outnumbered IgM particles, making IgM particle identification challenging. After iterative 2D classification, an IgM-enriched subset was subjected to ab initio reconstruction, heterogeneous refinement, and homogeneous refinement, yielding a low-resolution IgM reconstruction from ~19K particles. A 50-fold symmetry expansion was applied to the ~19K particles using the low-resolution IgM reconstruction as a reference, generating ~958K symmetry-expanded duplicate particles for particle "fishing." This expanded particle set yielded improved 2D class averages, facilitating more accurate particle matching and enabling recovery of IgM particles that had previously been mixed with other 2D classes. Following duplicate removal and additional 2D classification, one round of symmetry expansion-assisted particle fishing recovered ~183K IgM-like particles.

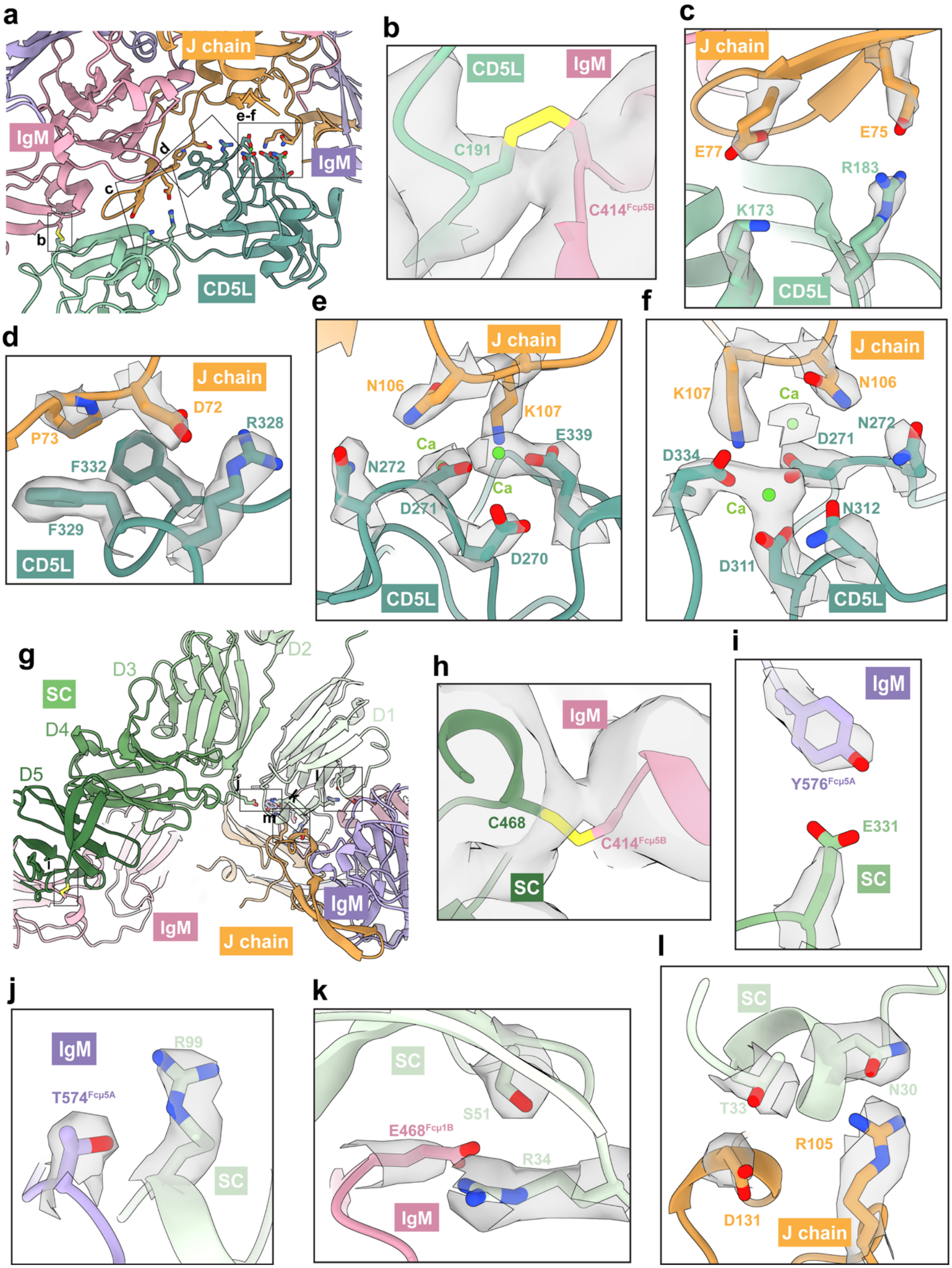
a
J chain
IgM
e-f
d
IgM
c
b
CD5L
b
CD5L
IgM
C191
C414Fcμ5B
c
J chain
E75
E77
R183
K173
CD5L
d
J chain
D72
P73
R328
F332
F329
CD5L
e
J chain
N106
K107
Ca
E339
N272
Ca
D271
D270
CD5L
f
J chain
K107
N106
Ca
N272
D271
D334
Ca
N312
D311
CD5L
g
D2
SC
D3
D1
D4
D5
j
k
l
m
i
IgM
IgM
J chain
h
IgM
C468
C414Fcμ5B
SC
i
IgM
Y576Fcμ5A
E331
SC
l
SC
N30
T33
R105
D131
J chain
j
IgM
R99
T574Fcμ5A
SC
k
SC
S51
E468Fcμ1B
R34
IgM

**Supplementary Fig. 6. Molecular interactions of CD5L and SC with the J chain and IgM-Fc core. a,** Overview of the CD5L-binding interface within the cavity of IgM-J. The regions shown in panels **b-f** are indicated. **b,** Disulfide bond between C191$^{CD5L}$ and C414$^{Fcμ5B}$. **c,** Polar interactions between the CD5L SRCR2 domain and the J chain, involving K173$^{CD5L}$-E77$^{J}$ and R183$^{CD5L}$-E75$^{J}$. **d,** Interactions between the CD5L SRCR3 domain and the J chain. F329$^{CD5L}$ and F332$^{CD5L}$ interact with P73$^{J}$, whereas R328$^{CD5L}$ forms a polar interaction with D72$^{J}$. **e-f,** $Ca^{2+}$-coordinating interactions at the CD5L-J-chain interface. **e,** Coordination of one $Ca^{2+}$ ion by D270$^{CD5L}$, D271$^{CD5L}$, E339$^{CD5L}$, N106$^{J}$, and K107$^{J}$, with an adjacent polar interaction between N272$^{CD5L}$ and N106$^{J}$. **f,** Coordination of a second $Ca^{2+}$ ion by D271$^{CD5L}$, D311$^{CD5L}$, N312$^{CD5L}$, D334$^{CD5L}$, N106$^{J}$, and K107$^{J}$, with the N272$^{CD5L}$-N106$^{J}$ polar interaction also shown. **g,** Overview of the SC-binding interface within the cavity of IgM-J. The regions shown in panels **h-l** are indicated. **h,** Disulfide bond between C468$^{SC}$ and C414$^{Fcμ5B}$. **i,** Polar interaction between the SC D3 domain and Fcμ5A, formed by E331$^{SC}$-Y576$^{Fcμ5A}$. **j,** Polar interaction between the SC D1 domain and Fcμ5A, formed by R99$^{SC}$-T574$^{Fcμ5A}$. **k,** Polar interactions between the SC D1 domain and the IgM Fcμ1B, involving S51$^{SC}$-E468$^{Fcμ1B}$ and R34$^{SC}$-E468$^{Fcμ1B}$. **l,** Polar interactions between the SC D1 domain and the J chain, involving T33$^{SC}$-D131$^{J}$ and N30$^{SC}$-R105$^{J}$.The color scheme is consistent with that used in **Fig. 2**. Cryo-EM densities are shown for the side chains depicted in **b-f and h-l**.

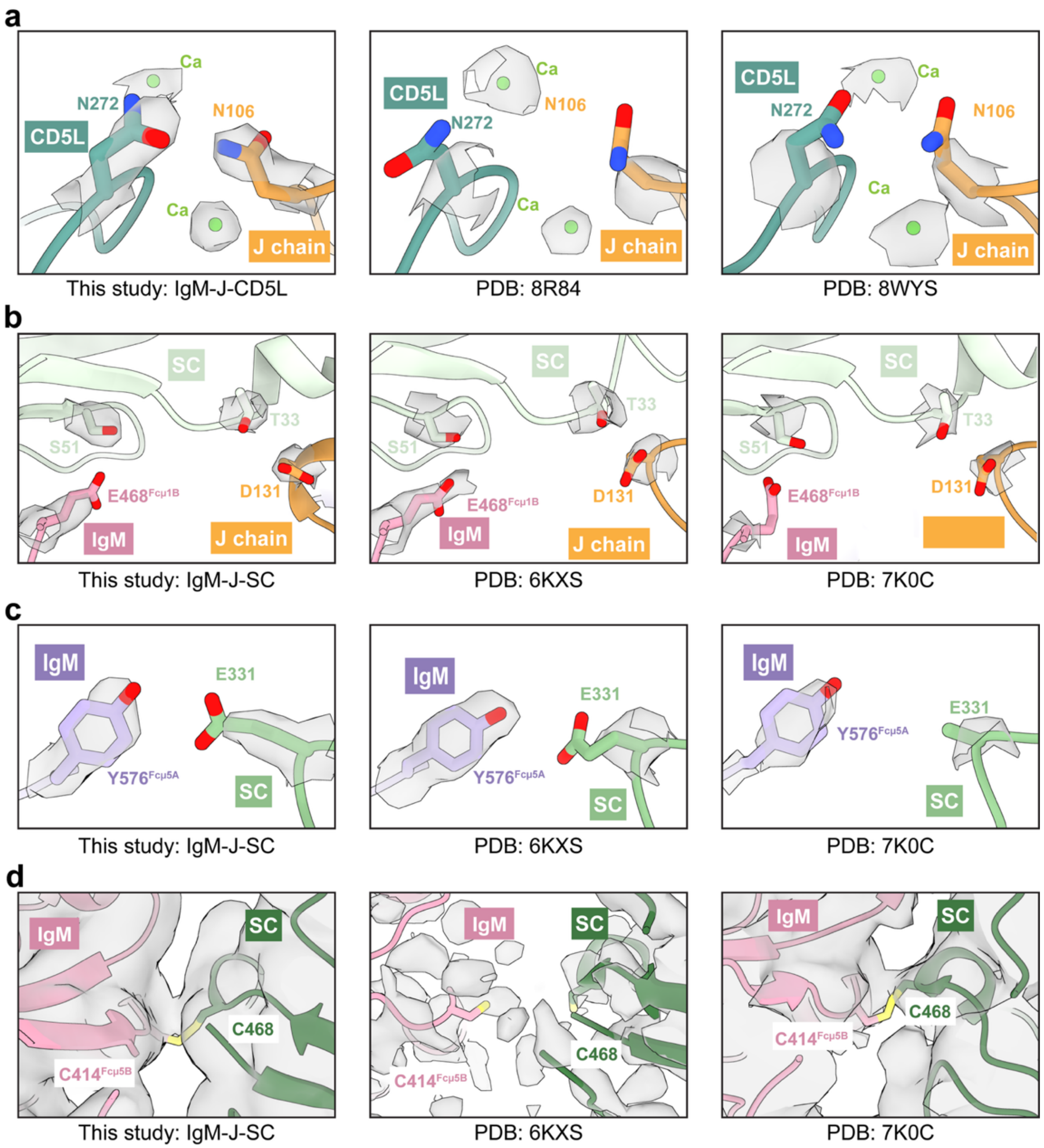


**Supplementary Fig. 7. Structural comparison of interaction sites in the IgM-J-CD5L and IgM-J-SC assemblies with previously published structures. a,** Comparison of the Ca$^{2+}$-binding pocket at the CD5L-J chain interface, highlighting the N106$^{J}$-N272$^{CD5L}$ interaction in the IgM-J-CD5L structure determined in this study (left) relative to previously deposited structures PDB 8R84 (middle) and PDB 8WYS (right). **b,** Comparison of the SC-IgM interface, highlighting the E331$^{SC}$-Y576$^{Fc\mu5A}$ interaction in the IgM-J-SC structure determined in this study (left) relative to PDB 6KXS (middle) and PDB 7K0C (right). **c,** Comparison of the SC-IgM and SC-J chain interfaces, highlighting the S51$^{SC}$-E468$^{Fc\mu1B}$ and T33$^{SC}$-D131$^{J}$ interactions in the IgM-J-SC structure determined in this study (left) relative to PDB 6KXS (middle) and PDB 7K0C (right). **d,** Comparison of the SC-IgM interface, showing the C414$^{Fc\mu5B}$-C468$^{SC}$ disulfide bond in the IgM-J-SC structure determined in this study (left) and PDB 7K0C (right), but not in PDB 6KXS (middle). The color scheme is consistent with that used in **Fig. 2**. Cryo-EM densities are shown for the side chains depicted.

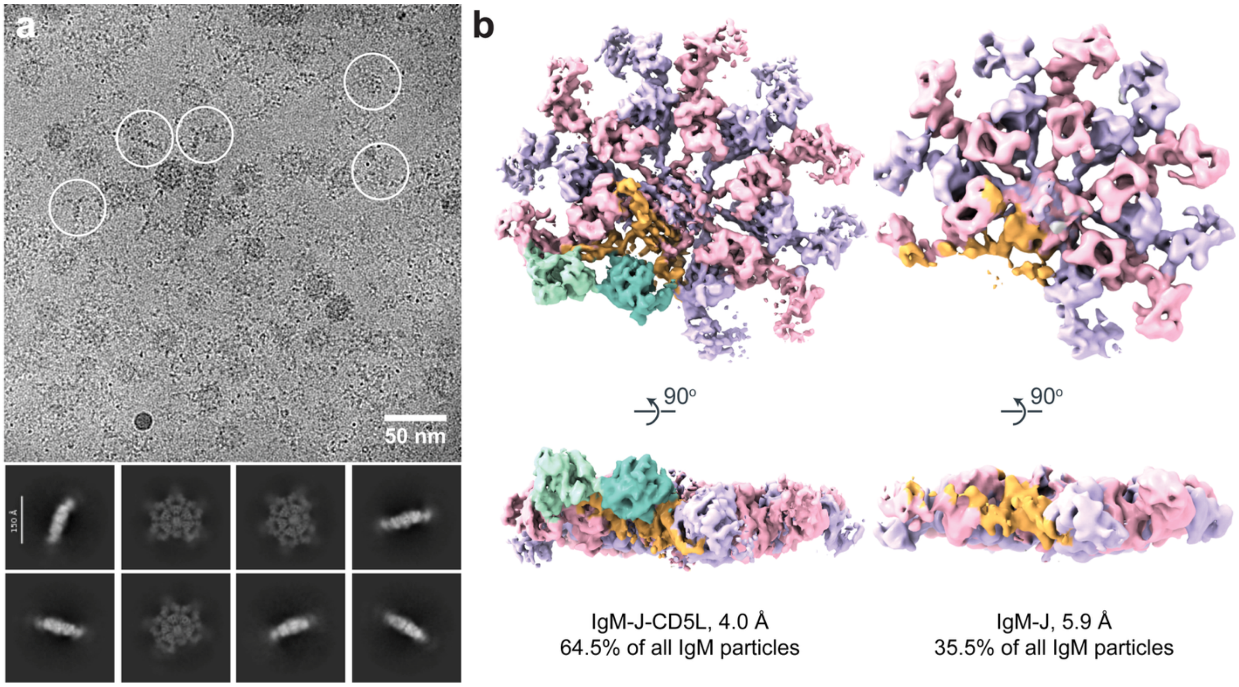


**Supplementary Fig. 8. Cryo-EM analysis of IgM complexes from a chronic patient serum sample. a,** Representative cryo-EM micrograph showing IgM and HBV SVPs, with selected IgM particles indicated by white circles. Representative top and side views of IgM 2D class averages are shown below. **b,** Top and side views of IgM-J-CD5L and IgM-J structures, with corresponding resolutions and particle-class distributions indicated. The color scheme is the same as **Fig. 2**.

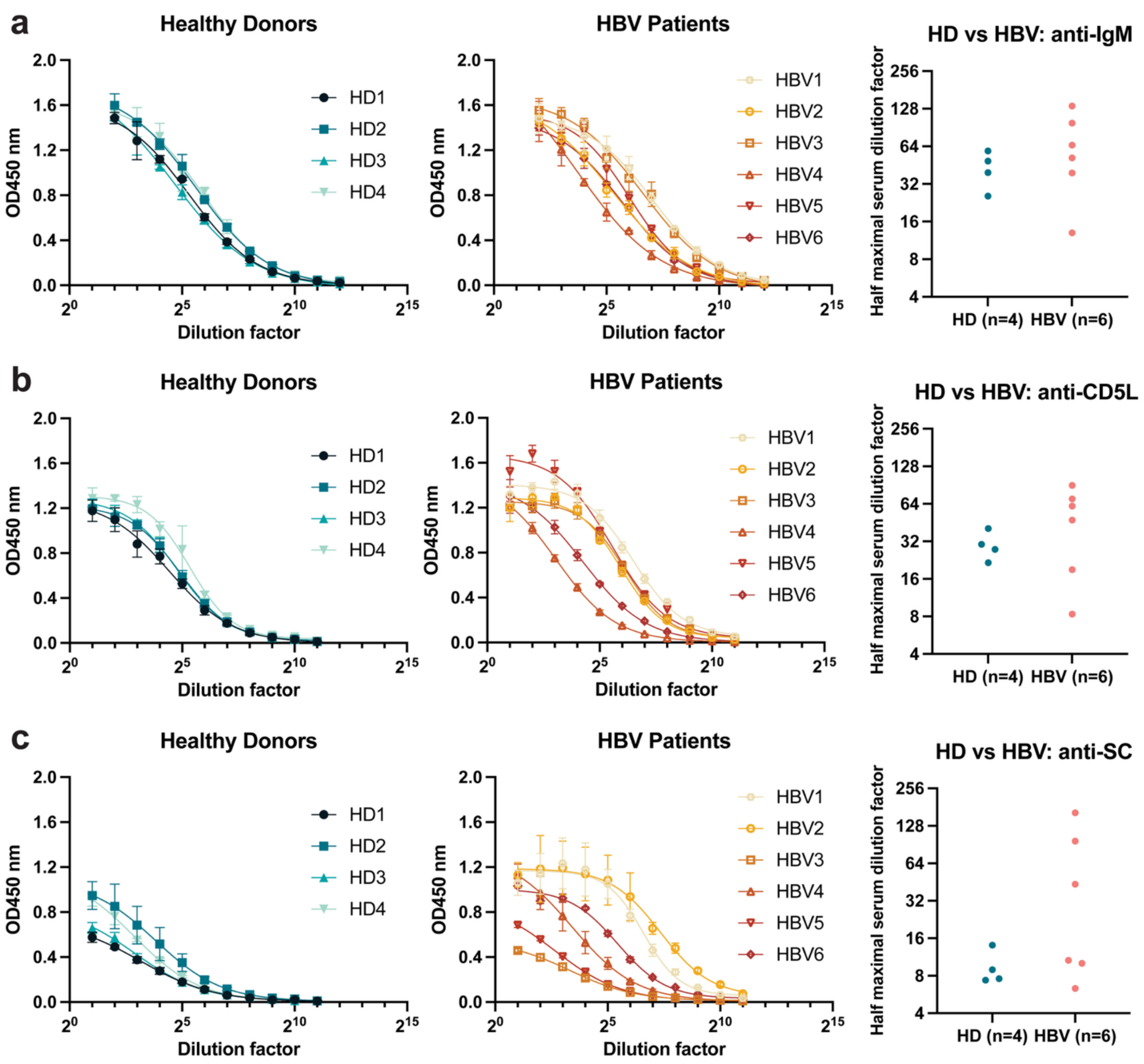


**Supplementary Fig. 9. ELISA analysis of serum IgM populations in healthy donors and HBV patients.** Serum samples from healthy donors (HD, n = 4) and HBV patients (n = 6) were serially diluted and analyzed by capture ELISA to assess total IgM and IgM associated with CD5L or SC. Plates were coated with antibodies against IgM (**a**), CD5L (**b**), or SC (**c**) to assess total IgM, CD5L-associated IgM, and SC-associated IgM, respectively. For each panel, ELISA dilution curves are shown for healthy donors on the left and HBV patients in the middle. Data are shown as mean ± SD of three technical replicates. The corresponding half-maximal serum dilution factors are shown for individual samples (right).

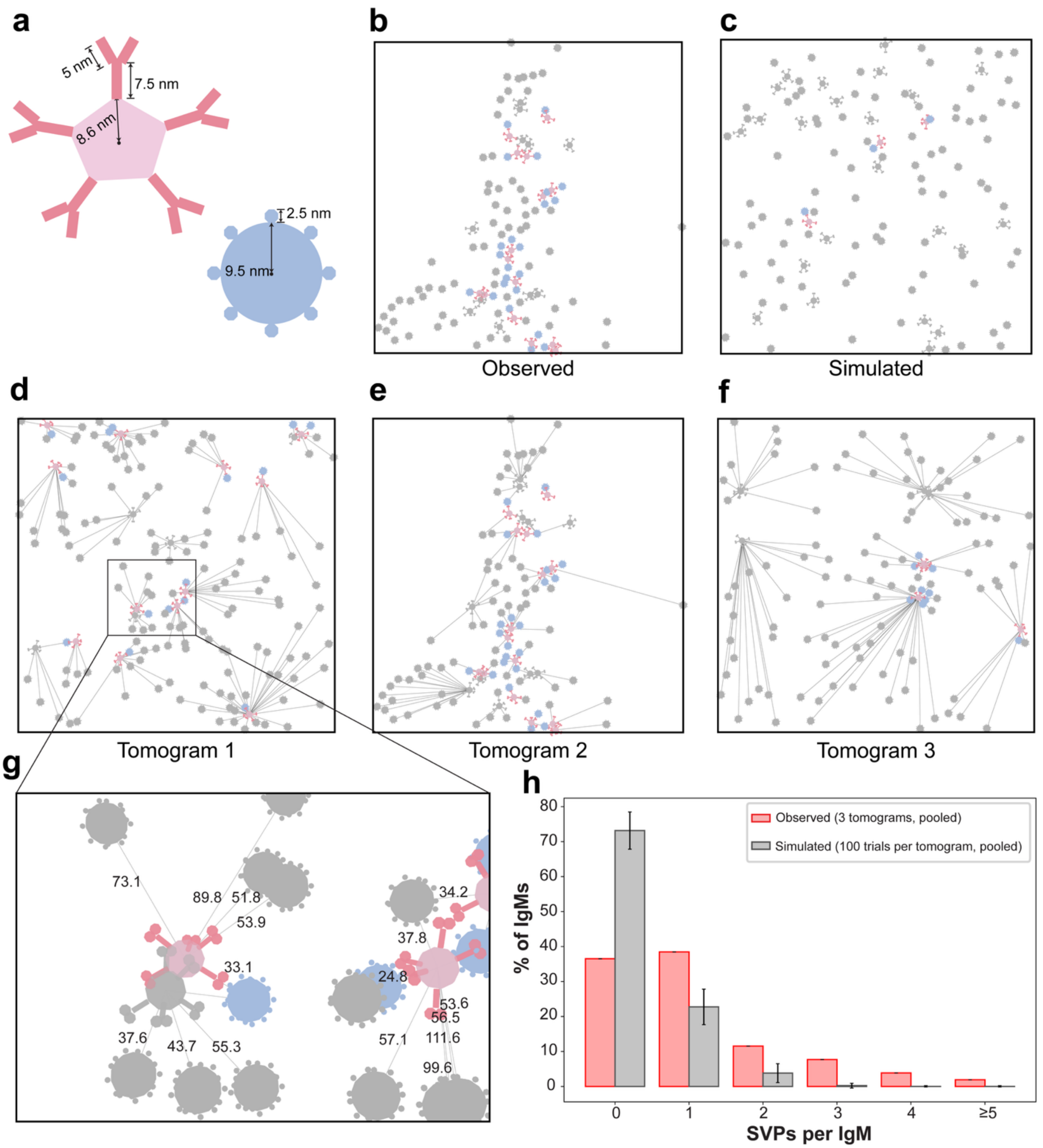

a
5 nm
7.5 nm
8.6 nm
2.5 nm
9.5 nm
b
Observed
c
Simulated
d
Tomogram 1
e
Tomogram 2
f
Tomogram 3
g
73.1
89.8
51.8
53.9
33.1
37.6
43.7
55.3
34.2
37.8
24.8
53.6
56.5
57.1
111.6
99.6
h
Observed (3 tomograms, pooled)
Simulated (100 trials per tomogram, pooled)
% of IgMs
SVPs per IgM
0
1
2
3
4
≥5

**Supplementary Fig. 10. Spatial simulation of IgM-SVP interactions. a,** Schematic representations of IgM and HBV-SVP particles, showing the geometric parameters used in the spatial simulations. **b-c,** Observed particle distribution in a real tomogram (**b**) and a simulated particle distribution using the dimensions and numbers of IgM and SVP particles from the real tomogram (**c**). **d-f,** Nearest-neighbor network map for tomograms 1 (**d**), 2 (**e**), and 3 (**f**), showing each SVP connected to its nearest IgM molecule. IgM-SVP pairs with center-to-center distances of less than 33.1 nm are indicated as interactions. All the IgM molecules within 40 nm radius of a given SVP are mapped to indicate multiple IgM molecules can interact with one SVP. **g,** Enlarged view of the boxed region in tomogram 1 illustrating center-to-center distance measurements between IgM and SVPs. Numbers indicate distances in nanometers. **h,** Distribution of the number of SVPs associated with each IgM molecule in the observed tomograms (red; three tomograms pooled) compared with spatially randomized simulations (grey; 100 simulations per tomogram, pooled). The mean numbers of SVPs per IgM were 1.093 for the observed tomograms and 0.323 for the simulations ($\sigma$ = 0.081).

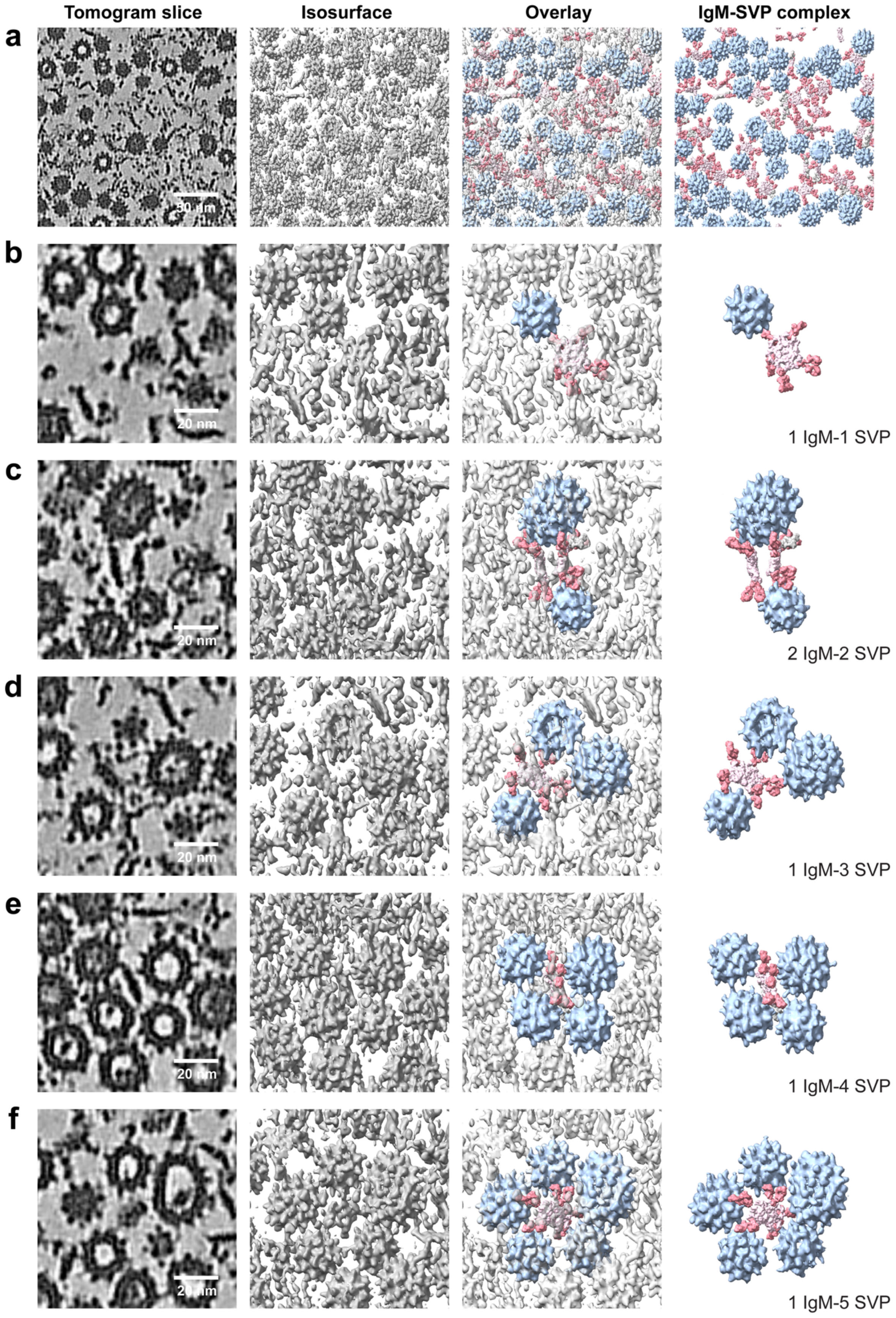

Tomogram slice
Isosurface
Overlay
IgM-SVP complex
a
30 nm
b
20 nm
1 IgM-1 SVP
c
20 nm
2 IgM-2 SVP
d
20 nm
1 IgM-3 SVP
e
20 nm
1 IgM-4 SVP
f
20 nm
1 IgM-5 SVP

**Supplementary Fig. 11. Cryo-ET reveals multivalent IgM-SVP interactions from an IgM-centered perspective. a,** Representative tomogram providing an overview of IgM and HBV SVPs, shown as a tomographic slice, corresponding isosurface rendering, overlay, and segmented IgM-SVP complexes from left to right. **b-f,** Representative IgM-SVP complexes illustrating different patterns of multivalent engagement, in which individual IgM molecules contact one to five SVPs. For each assembly, the tomographic slice, corresponding isosurface rendering, overlay, and segmented IgM-SVP complex are shown from left to right. Segmentation colors: SVPs, light blue; IgM Fc regions, light pink; Fab arms, salmon.

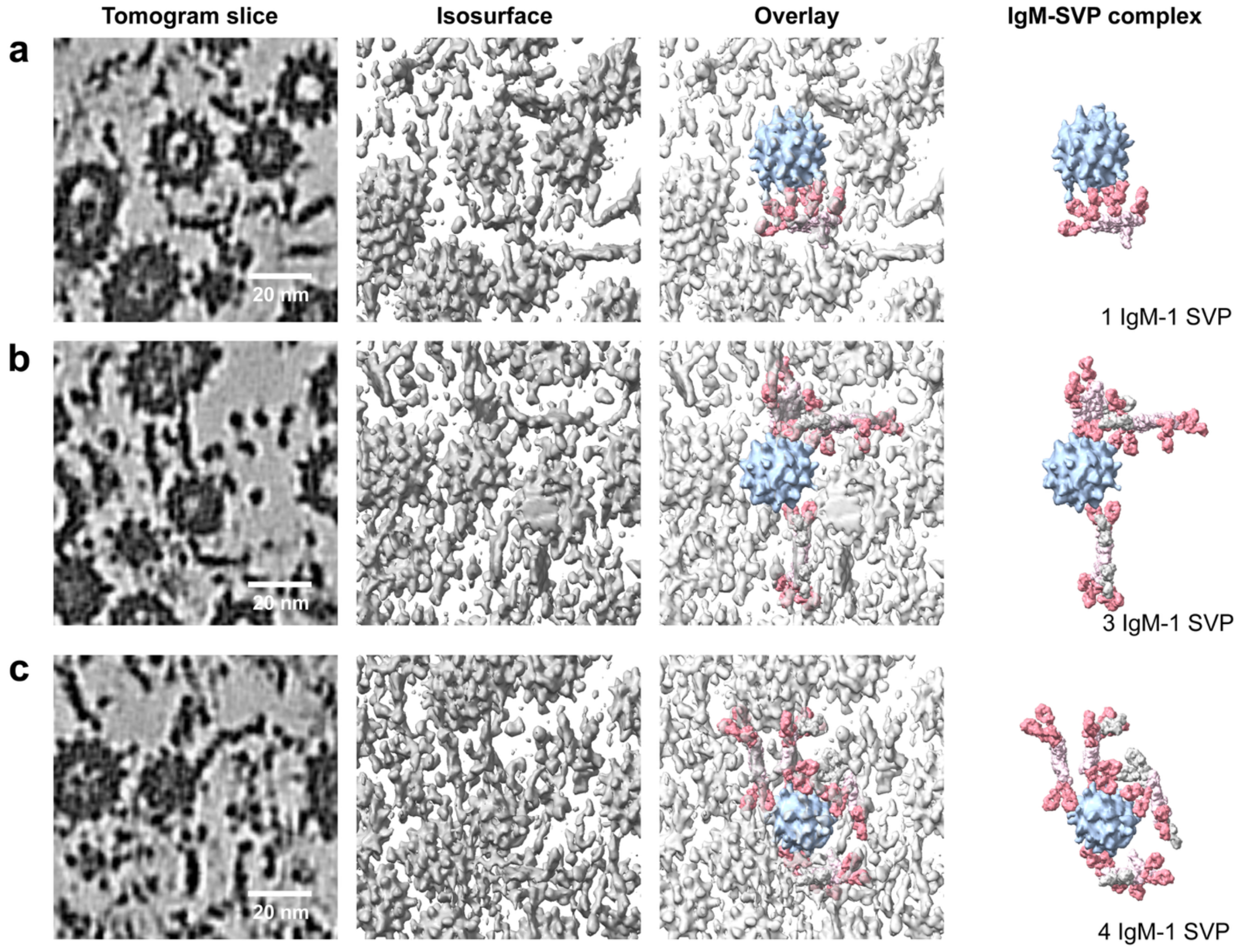


**Supplementary Fig. 12. Cryo-ET reveals multivalent IgM-SVP interactions from an SVP-centered perspective. a-c,** Representative IgM-SVP assemblies illustrating different patterns of multivalent engagement, in which individual SVPs contact one to four IgM molecules. For each complex, the tomographic slice, corresponding isosurface rendering, overlay, and segmented IgM-SVP complex are shown from left to right. Segmentation colors: SVPs, light blue; IgM Fc regions, light pink; Fab arms, salmon.

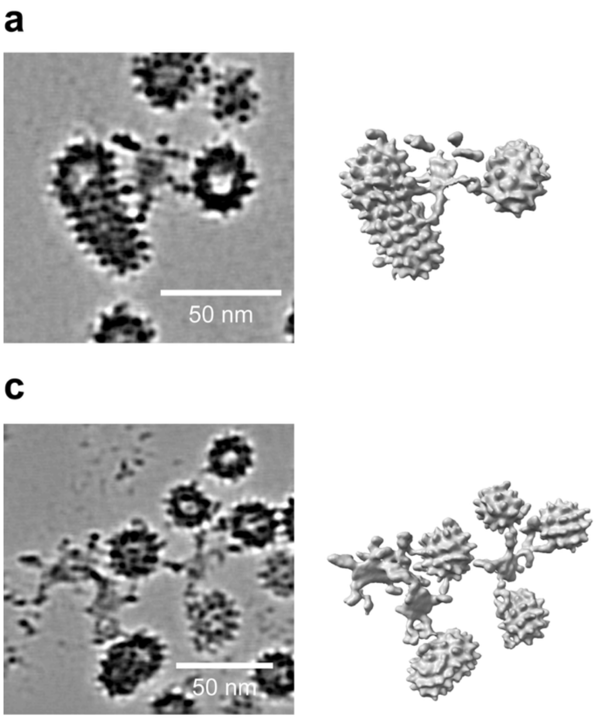


**Supplementary Fig. 13. IgM-SVP interactions persist at low concentration. a-b,** Representative tomographic slices and corresponding isosurface renderings showing IgM-SVP interactions in the low-concentration sample. **c-d,** Representative tomographic slices and corresponding isosurface renderings showing higher-order IgM-SVP immune complexes in the low-concentration sample.

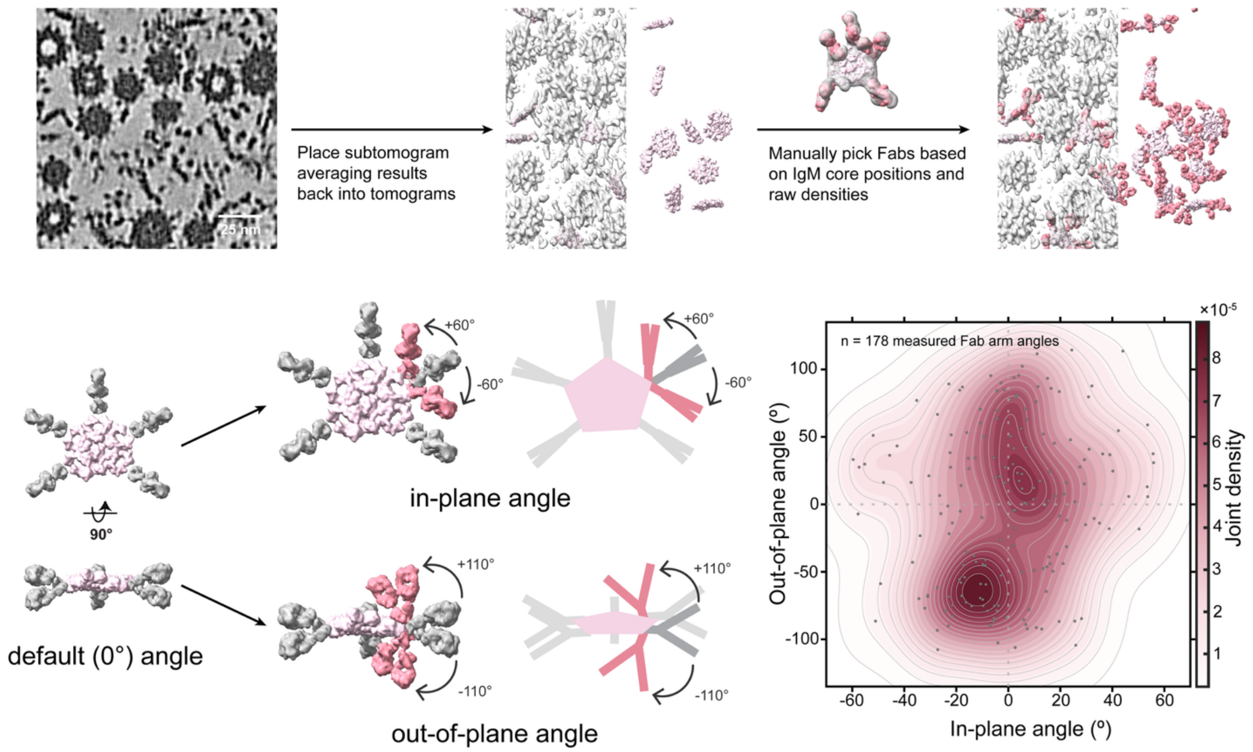


**Supplementary Fig. 14. Cryo-ET analysis of IgM Fab conformational space.** IgM Fc cores were placed back into tomograms improved by MissAlignment and IsoNet2 based on subtomogram averaging 3D refinement results, and flexible Fab arms were manually segmented from the corresponding tomographic densities. Fab arm orientations were determined by measuring the deviation of manually annotated Fab arms (salmon) from a default reference orientation (grey), and the resulting in-plane and out-of-plane distributions were used to define the reachable space of IgM Fabs. The joint distribution of the measured in-plane and out-of-plane angles from n = 178 manually annotated Fab arms is shown as a two-dimensional kernel density estimate (KDE), defining the experimentally observed conformational space of IgM Fab arms.

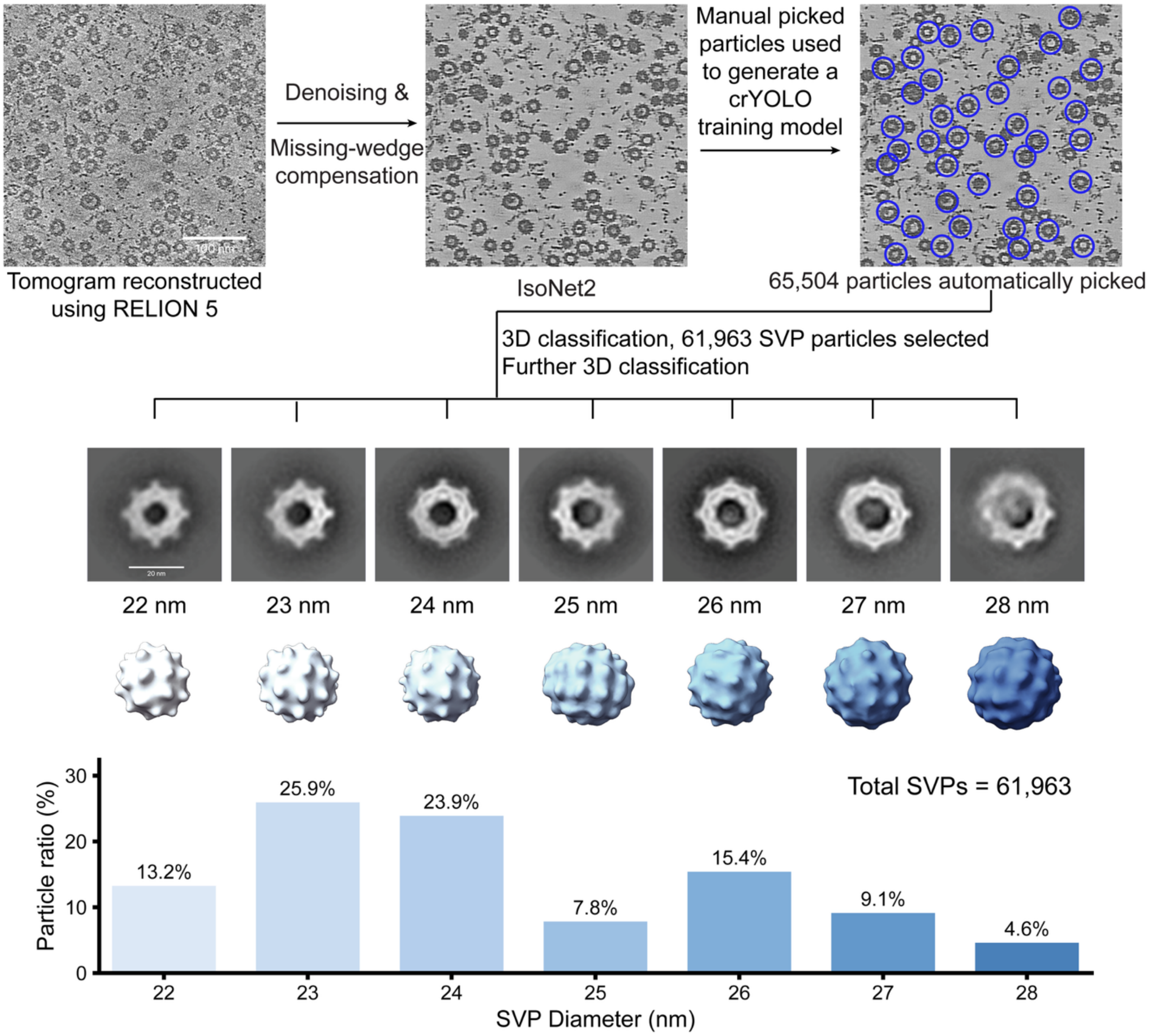


**Supplementary Fig. 15. Cryo-ET data-processing workflow for HBV SVP analysis.** Representative tomographic slice from a tomogram reconstructed using RELION 5. IsoNet2 was applied for denoising and missing-wedge compensation. Manually picked 1,051 HBV SVPs from 4 tomograms were used to generate a training model for crYOLO, followed by automated picking of 65,504 particles. After 3D classification, 61,963 SVP particles were selected and subjected to further 3D classification, revealing SVP populations with distinct diameters ranging from 22 to 28 nm. The relative abundance of each SVP population is shown at the bottom.

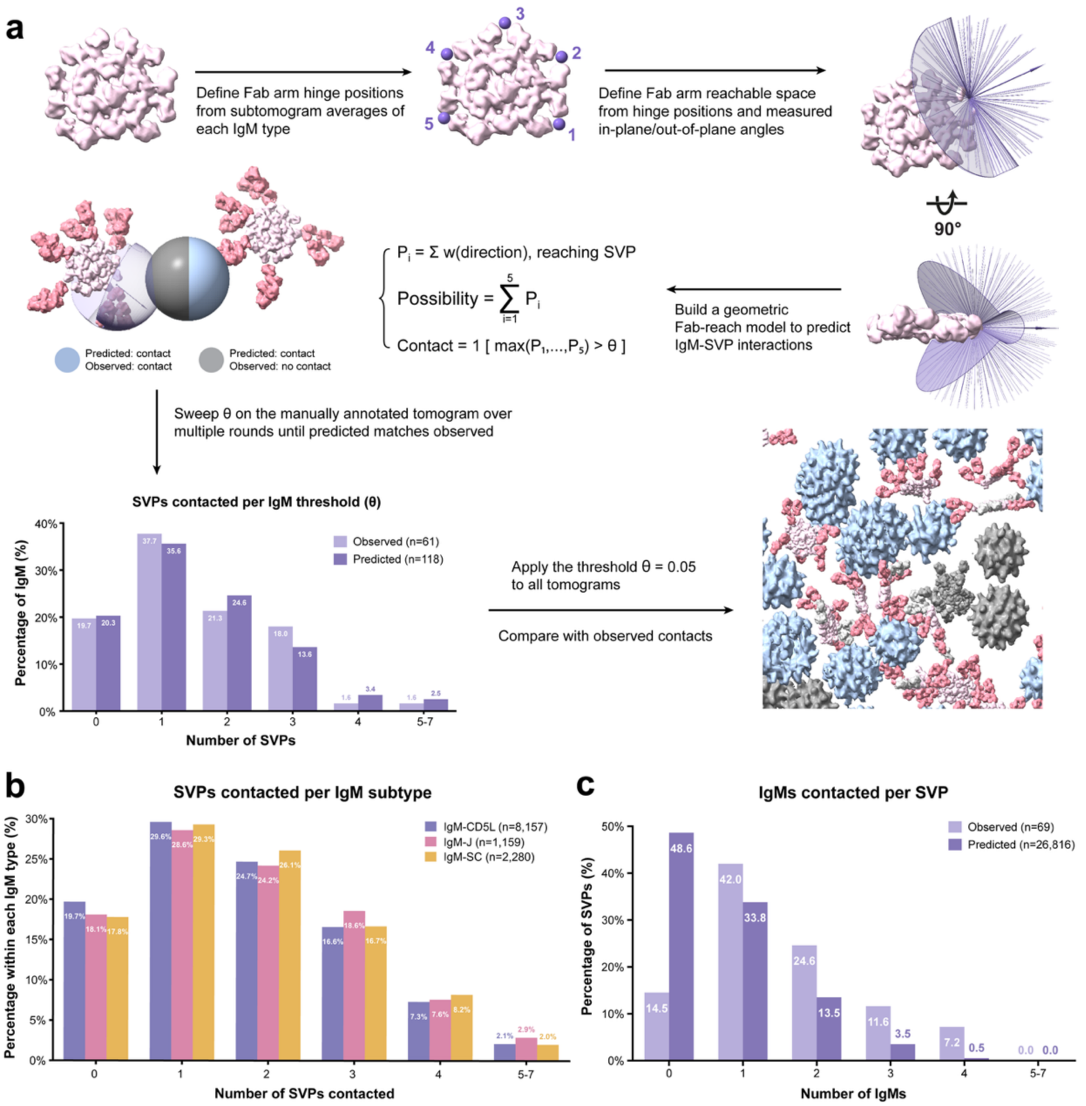


**Supplementary Fig. 16. Geometry-based prediction of IgM-SVP interactions. a,** Workflow for geometry-based prediction of IgM-SVP contacts. Fab hinges (purple points) were defined based on the STA-derived IgM core positions, and the experimentally determined Fab conformational space was used to define the spatial region reachable by each Fab arm. STA-derived HBV SVP positions and diameters were incorporated into the geometric model to predict potential IgM-SVP interactions. For each IgM-SVP pair, the probability of contact was calculated from the fraction of Fab conformations capable of reaching the SVP, and the probabilities across the five Fab arms were combined to estimate the overall contact probability. A probability threshold of θ = 0.05 was used to classify predicted contacts. The resulting distribution of SVPs contacted per IgM molecule was compared with manually annotated observations. Representative results are shown in segmented tomograms, with interacting particles colored and isolated particles shown in grey. **b,** Distribution of the number of SVPs contacted per IgM molecule, stratified by IgM subtype (IgM-J-CD5L, IgM-J, and IgM-J-SC). **c,** Comparison of the observed and geometry-based predicted distributions of the number of IgM molecules contacted per SVP.

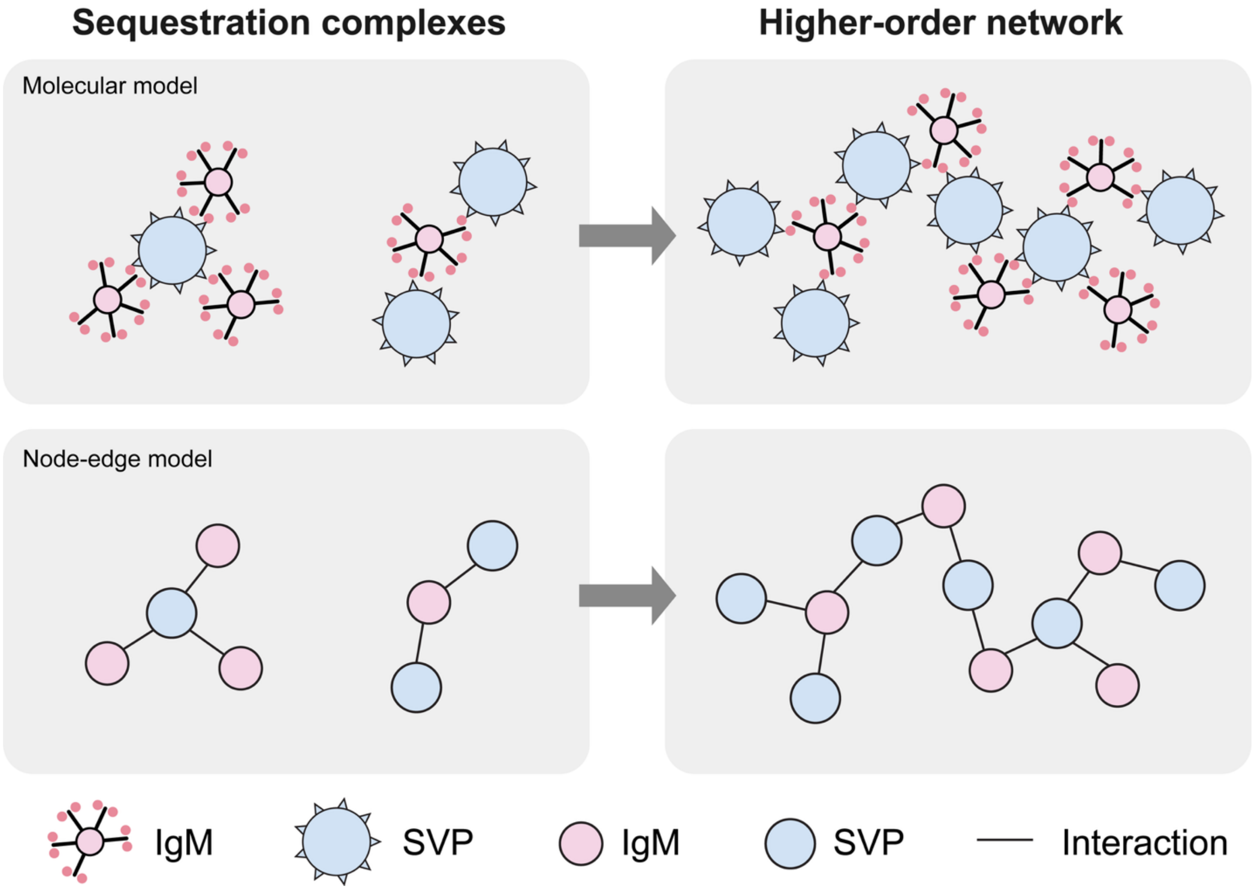


**Supplementary Fig. 17. Proposed model for IgM and SVP interaction.** Model for the transition from a passive sequestration mechanism, in which IgM and SVPs associate without interconnection, to a supramolecular immune complex mechanism, in which IgM crosslinks SVPs into an interconnected higher-order network.

### Supplementary Movies

**Supplementary Movie 1.** In situ visualization of IgM-SVP interactions by cryo-ET. Visualization of segmented SVPs and full-length IgM structures within the tomogram, followed by a close-up view highlighting the IgM-SVP interactions.

**Supplementary Movie 2.** IgM-SVP interactions at a lower sample concentration. Visualization of IgM-SVP complexes alongside isolated IgM molecules and SVPs, supporting that the observed interactions are not artifacts induced by high sample concentration.

**Supplementary Movie 3.** Molecular interactions of CD5L and SC with IgM-J. Visualization of the CD5L- and SC-binding interfaces with IgM-J, highlighting the newly resolved interactions identified in this study.

**Supplementary Table 1. Clinical information for patients included in this study.**

| | **Patient 1** | **Patient 2** |
|---|---|---|
| **Study inclusion date** | March 21, 2024 | September 19, 2024 |
| **Lab ID** | P0321 | P1017 |
| **Sex** | Male | Male |
| **Date of birth** | April 9, 1940 | January 1, 1988 |
| **Main diagnosis** | Died on March 22, 2024, due to acute Hepatitis B virus (re-)infection | Chronic HBV |
| **HBV exposure/diagnosis date** | n/a | 09/23 |
| **HBV genotype** | D1 | D1 |
| **HBsAG quant IU/mL** | 37,325 | 30,120 |
| **HBV DNA abs. IU/mL** | 109,000,000 | 212,000 |
| **GOT/AST** | 395 | 16 |
| **GPT/ALT** | 772 | 35 |
| **Fibroscan (E) kPa** | 12.6 | 6 |
| **HBeAg Date** | n/a | November 16, 2023 |

**Supplementary Table 2. Characteristics of HBV-positive donor sera used for ELISA.**

| Sample | Sample ID | Collection date | HBV viral load (IU/mL) | Viral-load test date |
|---|---|---|---|---|
| HBV1 | 533876-10 | October 1, 2025 | 279,510,695 | December 11, 2025 |
| HBV2 | 533848-10 | October 1, 2025 | 324,410,193 | December 11, 2025 |
| HBV3 | 441619-12 | February 23, 2023 | 490,978,867 | March 23, 2023 |
| HBV4 | 533883-16 | October 1, 2025 | 757,729,537 | December 11, 2025 |
| HBV5 | 437842-29 | January 25, 2023 | 404,273,969 | February 23, 2023 |
| HBV6 | 533931-14 | October 1, 2025 | 614,928,730 | December 11, 2025 |

**Supplementary Table 3. Cryo-ET data acquisition and image processing.**

| **Sample (cryo-ET)** | **IgM assemblies** | | |
|---|---|---|---|
| Microscope | TFS Titan Krios G3 | | |
| Voltage (keV) | 300 | | |
| Energy-filter (eV) | 10 | | |
| Detector | Falcon 4i | | |
| Magnification | 81,000 | | |
| Pixel size (Å) | 1.512 | | |
| Defocus range (μm) | -2 to -6 | | |
| Acquisition scheme | Dose-symmetric, −60°/+60°, 3° | | |
| Total Dose (electrons/Å$^2$) | ~120 | | |
| Tomograms (used / collected) | 253/269 | | |
| Total particles (used / picked) | 18,368/95,302 | | |
| **IgM assemblies** | **IgM-J-CD5L** | **IgM-J** | **IgM-J-SC** |
| Symmetry | C1 | C1 | C1 |
| Final particles | 18,359 | 2,744 | 4,921 |
| Resolution at 0.143 FSC (Å) | 7.6 | 11.4 | 8.8 |

**Supplementary Table 4. Statistics of cryo-EM data collection and image processing.**

| Sample (cryo-EM) | IgM assemblies | | |
|---|---|---|---|
| Microscope | TFS Titan Krios G3 | | |
| Voltage (keV) | 300 | | |
| Spherical aberration (mm) | 2.7 | | |
| Energy-filter (eV) | 10 | | |
| Detector | Falcon 4i | | |
| Magnification | 105,000 | | |
| Pixel size (Å) | 0.93 | | |
| Defocus range (μm) | -0.7 to -2.4 | | |
| Total Dose (electrons/Å$^2$) | 50 | | |
| Software for data collection | EPU | | |
| Number of micrographs | 75,331 | | |
| **IgM assemblies** | **IgM-J-CD5L** | **IgM-J** | **IgM-J-SC** |
| Symmetry imposed | C1 | C1 | C1 |
| Box size (pixel) | 384 | 384 | 384 |
| Final particle images (no.) | 557,671 | 223,874 | 260,476 |
| Resolution at FSC=0.143 (Å) | 2.81 | 3.01 | 3.05 |

**Supplementary Table 5. Statistics of cryo-EM model refinement.**

| | **IgM-J-CD5L** | **IgM-J** | **IgM-J-SC** |
|---|---|---|---|
| Model resolution at FSC=0.5 (Å) | 2.9 | 3.2 | 3.1 |
| Model compositions | | | |
| Chains | 12 | 11 | 12 |
| Atoms | 40650 (Hydrogens: 20079) | 37304 (Hydrogens: 18462) | 45285 (Hydrogens: 22398) |
| Residues | Protein: 2611 | Protein: 2390 | Protein: 2901 |
| Ligands | NAG: 18; Ca: 2 | NAG: 17 | NAG: 23 |
| Bonds (RMSD) | | | |
| Length (Å) (# > 4σ) | 0.005 (0) | 0.004 (0) | 0.004 (0) |
| Angles (°) (# > 4σ) | 1.012 (0) | 1.063 (0) | 1.003 (0) |
| MolProbity score | 1.03 | 1.11 | 1.20 |
| Clash score | 2.44 | 3.22 | 4.20 |
| Ramachandran plot (%) | | | |
| Outliers | 0.00 | 0.00 | 0.00 |
| Allowed | 1.59 | 1.14 | 1.78 |
| Favored | 98.41 | 98.86 | 98.22 |
| Rotamer outliers (%) | 0.64 | 0.23 | 0.62 |
| Cβ outliers (%) | NA | NA | NA |
| Peptide plane (%) | | | |
| Cis proline/general | 5.1/0.0 | 5.3/0.0 | 4.7/0.0 |
| Twisted proline/general | 0.0/0.0 | 0.0/0.0 | 0.0/0.0 |
| CaBLAM outliers (%) | 1.41 | 0.85 | 1.66 |
| ADP (B-factors) | | | |
| Iso/Aniso (#) | 20571/0 | 18842/0 | 22887/0 |
| min/max/mean | | | |
| Protein | 8.61/186.46/90.95 | 1.91/186.27/82.89 | 4.93/209.42/105.48 |
| Ligand | 39.06/86.23/63.93 | 40.54/112.31/72.87 | 35.13/160.24/85.33 |
| EMRinger score | 3.11 | 2.84 | 3.08 |
